# Tetralone-ABA enhances winter cold acclimation, reduces deacclimation, and delays budbreak in V. vinifera and V. hybrid grapevines

**DOI:** 10.1101/2024.12.31.630556

**Authors:** Hongrui Wang, Yue Pan, Jason P. Londo

**Author notes:** Correspondence: Corresponding Authors Hongrui Wang, Jason Londo (jpl275@cornell).

## Abstract

Climate change-related acute winter freezes and unseasonal false springs have become significant and predictable risks for grape growers across North America and Eurasia. Novel strategies to enhance resilience during the dormant season, particularly by improving bud cold hardiness and delaying budbreak, are urgently needed for the sustainability of grape and wine production. In this study, we evaluated a synthetic abscisic acid (ABA) analog, tetralone-ABA (ABA-1102), as a sprayable product for inducing these traits in three grapevine cultivars: ‘Riesling’, ‘Cayuga White’, and their progeny, ‘Aravelle’. We determined that post-harvest foliar application of tetralone-ABA promoted early leaf senescence, induced cultivar-specific bud cold hardiness enhancement during cold acclimation, slowed deacclimation under controlled and field conditions, and delayed budbreak in spring without affecting growing season phenology, physiology or harvest yield. Transcriptomic analysis of during deacclimation assays suggests that tetralone-ABA’s effect on delaying deacclimation may result from its suppression of the activation of growth-related pathways under growth-permissive conditions. Detailed investigations of these pathways indicate that tetralone-ABA may have modulated critical biological processes such as cell wall remodeling, sugar metabolism, and ABA signaling. Overall, this study provides novel insights into the genetic control of grapevine deacclimation, highlights ABA’s role in grapevine dormant season physiology, and demonstrates tetralone-ABA’s potential as a promising tool for improving dormant season viticulture resilience, offering a new strategy to protect grapevines against the increasing threats posed by climate change.

## 1. Introduction

The growing frequency and intensity of extreme weather events due to climate change are emerging as the primary obstacles to sustainable agriculture and global food security [1–3]. Climate-related extreme meteorological events, hydrological events and climatological events threaten the productivity of the major agricultural systems. Perennial crops, such as grapevine, could be more vulnerable to these events due to long lifespan and high re-establishment cost for the adoption of new cultivars [4]. Among all the abiotic stresses associated with climate change, cold stress during the dormant season is a leading factor constraining the expansion of the grape and wine industries in cool climate viticulture regions in North America and Eurasia [5–9]. Low temperatures in winter can exceed the cold hardiness of grapevines, resulting in tissue damage to the compound bud, phloem and xylem or the whole plant [6]. Cold damage decreases productivity in the following season, and the cost of replanting and additional managements in the following years intensively impact the profitability of the damaged vineyards [10,11]. Globally, the increasing frequency of late spring frost damage due to false spring (advanced budbreak in overall warming winter followed by late frost) is becoming a leading challenge in the major viticultural regions in the world [12]. Late spring frosts directly threaten the growth of grapevines, especially the most fruitful primary buds, leading to significant yield loss [7,13]. Thus, the enhancement of dormant season viticultural resilience, is crucial for the sustainability of global grape and wine industry.

Active preventative methods that directly increase vineyard ambient temperature or canopy temperature through wind machines, heaters, helicopters, overhead sprinklers or hydrophobic particle film and acrylic polymer have been applied in commercial vineyard to manage cold-related damage [7,12,14]. However, to date, these methods are not economically feasible for smaller acreage growers due to their high capital cost. To this end, researchers have been developing novel tools to mitigate cold-related damage, and the focuses are mainly on delaying budbreak to avoid frost damage in spring and enhancing cold hardiness to mitigate cold damage in winter. Pre-budbreak application of plant oil-based products such as Amigo Oil and FrostShield were determined to be effective in delaying budbreak from days to weeks [13,15,16]. However, some studies report some carry-on effects from such applications, like phytotoxic effects on certain cultivars contributing to lower bud survival rate and cluster number during harvest [16–18]. Delayed pruning and double pruning are other examples of approaches aiming to delay budbreak on basal buds through the prolonged apical dominance of apical buds. Such approaches have been shown effective in delaying budbreak for 15 to 20 days, but the delaying effect can last until harvest, which impacts fruit maturity and wine quality [12]. Apart from the side effects of these methods, the increased complexity of vineyard management may also slow the adaptation of these methods among growers. Methods that achieve budbreak delay while avoiding carry-on effects on grapevine growing and exhibit ease of application are needed to manage late spring frost damage in a changing world.

To mitigate winter cold damage in vineyards, grapevine breeders in cold/cool climate viticulture regions in the world have developed cold hardy cultivars by introgressing the superior cold hardiness of wild *Vitis* species adapted to cold winter conditions such as *V. labrusca* and *V. amurensis* into cold sensitive *V. vinifera* cultivars [19,20]. In parallel, the development of practical tools to enhance grapevine cold hardiness has centered on the evaluation of post-harvest application of plant hormones such as abscisic acid (ABA) [13,21–25]. ABA is an essential plant phytohormone that regulates the plant life cycle and functions in response to various abiotic stresses through the ABA signaling pathway and mediated by ABA responsive genes and ABA responsive element binding proteins (AREBs) [26]. The processes of dormancy establishment, transition, and release in perennials are controlled by the interaction of ABA and other plant hormones [27–29]. In grapevine, bud ABA concentration increases during cold acclimation (gain of cold hardiness) and decrease during deacclimation (loss of cold hardiness) as a result of altered ABA biosynthesis and catabolism during the dormant season, supporting ABA’s involvement in cold hardiness [30–32]. Application of ABA on grapevines upregulated the regulatory or functional pathways for cold hardiness such as C-Repeat Binding Factor/*DRE* Binding Factors (*CBF/DREB1*), starch, raffinose family oligosaccharides and hydrogen peroxide biosynthesis [13,25,33,34]. Some studies have reported that post-harvest foliar application of ABA on grapevine promoted cold acclimation, induced deeper dormancy, decrease bud water content and enhanced bud cold hardiness, but others reported limited effect [22,35–37]. These inconsistencies may be a result of natural ABA’s short half-life or natural rapid enzymatic catabolism as a phytotoxicity that interferes the equilibrium of transcriptome and metabolome [38]. In recent years, synthetic ABA analogs (8’-acetylene-ABA and tetralone-ABA) have been tested for their efficacy to enhance freezing tolerance in winter [36,39,40]. The analogs are functionally similar to the natural form of ABA but are structurally different, which contributes to increased resistance to catabolism [38]. Although previous studies reported that there were inconsistent or no effect during grapevine cold acclimation, post-harvest foliar application of ABA analogs significantly enhanced grapevine bud cold hardiness during deacclimation in late winter and significantly delayed budbreak for days in various cultivars [36,39,40]. Such products with relatively easy application methods and efficiency in budbreak delay might be of great interest among grapevine growers. However, a more in-depth evaluation of the impact of ABA analogs on grapevine physiology is needed.

In this study, we conducted a two-year evaluation of the impact of the post-harvest foliar application of an ABA analog, tetralone-ABA, on three grapevine cultivars through assessing changes in physiological, phenological and transcriptomic responses. We hypothesized that post-harvest foliar application of tetralone-ABA would enhance cold hardiness in winter and delay budbreak in spring, and the delaying effect is a consequence of tetralone-ABA-induced slower activation of growth resumption-related pathways. The first objective was to determine if tetralone-ABA could enhance grapevine bud cold hardiness and if the enhancements differ among genotypes. The second objective was to understand tetralone-ABA’s impact on the grapevine bud transcriptome during deacclimation under forcing conditions. The third objective was to evaluate if tetralone-ABA could delay budbreak, and if there were any fundamental changes in growing season physiology or harvest yield.

## 2. Materials and Methods

### 2.1 Plant materials and treatment application

The experiment was conducted at the Finger Lakes Teaching and Demonstration Vineyard at Anthony Road Wine Co. at Geneva, NY, US (42.71° N, −76.97 ° W). The experiment was conducted in two years: 2022-2023 and 2023-2024. Treatment applications occurred on 10/05/2022 (2022-2023 experiment) and 10/17/2023 (2023-2024 experiment) on *V. vinifera* ‘Riesling’, *V. hybrid* ‘Cayuga White’ and their progeny, ‘Aravelle’. ‘Riesling’ is a moderate cold hardiness cultivar of great economic importance for cool and cold climate grape and wine industries such as Finger Lake regions. ‘Riesling’ exhibits mid-season budbreak as compared to other *V. vinifera* cultivars [41]. ‘Cayuga White’, with *V. labrusca* and *V. rupestris* and *V. aestivalis* genetic background, sustains sufficient cold hardiness to survive in the winters in the cool climate viticultural regions in North America. This cultivar tends to deacclimate slower and initiate budbreak later as compared to other cold hybrid cultivars that derive from *V. riparia*. Limited information is available for the overwintering behavior of ‘Aravelle’ as this cultivar was recently released.

Experimental vines were subjected to standard vineyard management during the growing season. Treatments of either tetralone-ABA treatment or water control were applied as a foliar canopy spray a week after the harvest of ‘Riesling’ on 10/05/2022 (for the 2022-2023 experiment) and 10/17/2023 (for the 2023-2024 experiment). Tetralone-ABA ((+)-tetralone ABA, ABA-1102, purity = 100%, ABAzyne Biosciences Inc., Saskatoon, Canada) was obtained as a fine powder and was kept under dry, cool and dark conditions to maintain chemical activity [38]. Stock solutions were prepared by dissolving 6 g of tetralone-ABA in 1 L of distilled water with 120.0 mL of DMSO (Fisher Scientific, NH, US) and 3840.0 µL of 5.00 M NaOH solution (Sigma-Aldrich, MO, US). Stock solutions were diluted to the final concentration of 0.5 g·L^-1^ using distilled water. During treatment application, 1 L of 0.5 g·L^-1^ tetralone-ABA solution was applied on the vegetative canopy of each experimental vine, corresponding to a final application rate of 0.5 g tetralone-ABA per vine. Control treatment was distilled water mixed with the same concentrations of DMSO and NaOH. Silwet L-77 (Helena Agri-Enterprises, TN, US) was added at a rate of 0.05 % (v/v) to both treatments as a surfactant. For each cultivar, 12 individual vines were treated in both tetralone-ABA and control treatments. To avoid any carry-one effect from the treatment in the previous year, different vines were used for experimentation in each of the two dormant seasons.

### 2.2 Relative chlorophyll concentration measurement

In the 2022-2023 experiment, relative chlorophyll concentration of the leaves in experimental vines were quantified after treatment application using an MC-100 chlorophyll concentration meter (Apogee Electronics, CA, USA). For each vine, the measurement was conducted three times on three randomly selected leaves in the lower, middle, and upper canopy. Relative chlorophyll concentration was expressed as SPAD (Soil Plant Analysis Development), a relative measure of chlorophyll concentration derived from the absorbance of specific wavelengths of light [42–44]. The three readings were averaged as subsamples to represent the relative chlorophyll concentration of the vine. Measurements were conducted four times until defoliation at one-day post-treatment (10/06/2022), one-week post-treatment (10/12/2022), two-week post-treatment (10/19/2022) and three-week post-treatment (10/26/2022).

### 2.3 Cold hardiness monitoring

After the treatment application, the monitoring of grapevine bud cold hardiness was conducted over the entire dormant season to evaluate the treatment effect. Grapevine bud cold hardiness was determined bi-weekly from 10/26/2022 to 03/22/2023 in the 2022-2023 dormant season and from 10/17/2023 to 3/18/2024 in the 2023-2024 dormant season using differential thermal analysis based on a standardized protocol [9,45]. Briefly, grapevine buds were loaded in sample plates and treated with a decreasing temperature ramp of 4 °C·h^-1^ from 0 °C to −50 °C in a Tenney T2C environmental chamber (Tenney Environmental, PA, US). Thermocouples on samples plates detect low temperature exotherms (LTEs), which represent the intracellular ice nucleation of grapevine buds, corresponding to the cold hardiness of the buds [45,46]. For each cultivar and each treatment, at least 15 buds were used to determine the concurrent cold hardiness.

### 2.4 Deacclimation assays

In addition to bud cold hardiness monitoring, deacclimation rate, the speed of cold hardiness loss that occurs under growth permissive condition, was measured three times in early winter, mid-winter and late winter to monitor the progression of dormancy transition in response to tetralone-ABA treatment. Deacclimation rate was only determined in ‘Riesling’ due to limited plant materials for the other two cultivars. In the 2022-2023 experiment, buds were collected three times from the field at 518.5 (11/07/2022, early winter), 978.5 (01/02/2023, mid-winter) and 1339.9 (03/13/2023, late winter) chilling units (Utah model) [47]. In the 2023-2024 experiment, buds were collected three times from the field at 629.1 (11/08/2023, early winter), 1282.6 (01/24/2023, mid-winter) and 1737.9 (03/20/2024, late winter) chilling units (Utah model). These three timepoints roughly correspond to the major physiological stages in dormant season; endodormancy (unable to deacclimate and initiate budbreak under growth permissive condition), transition from endodormancy to ecodormancy (partially able to deacclimate and initiate budbreak under growth permissive condition) and ecodormancy (fully able to deacclimate and initiate budbreak under growth permissive condition) [48]. During each deacclimation assay, canes collected from field were chopped into single bud cuttings, and the cuttings were incubated in growth chamber at 22 °C without supplemental light with cutting ends dipped in water. On day 0, day 3 (day 2 in the 2023-2024 experiment), day 5, day 7 and day 14 in the growth chamber, bud cold hardiness was determined by DTA if the buds remained unbroken. Deacclimation rate, the coefficient of the linear regression between LTE and days in chamber, was computed for each deacclimation assay and for each treatment [49].

### 2.5 Spring phenology monitoring

Phenology recording started before budbreak to evaluate treatment effect on growth restoration and vegetative growth during the growing season. Given one of the major goals of the study is to assess the impact of tetralone-ABA on the timing of budbreak, we combined two previously established grapevine phenology systems, EL system [50] for the stages after budbreak and Modified Shaulis Field Score (https://lergp.com/modified-shaulis-field-score) for the stage before budbreak to increase the resolution of phenological stages in the early growing season. Visual assessment of phenology was conducted on each experimental vine three times every week from 04/19/2023 to 06/26/2023 in the 2023 growing season and 04/23/2024 to 06/03/2024 in the 2024 growing season. For each cultivar and each treatment, the date that at least 50% of experimental vines reached budbreak stage was recorded as the date of budbreak [13,37,51]. We first quantified the delay of budbreak for the two treatments each year using ‘days’. To increase the precision of delaying effect from tetralone-ABA treatment, we also quantified the delay in thermal unit of GDH4 (growing degree hour using 4 °C as based temperature). The measuring method of GDH4 utilizes two cosine functions between a base temperature (4 °C) and an upper limit temperature (36 °C), with optimum at 26 °C as suggested in previous modeling approaches of grapevine and tree fruit budbreak [52,53].

### 2.6 Growing season physiology and harvest data

During the growing season, grapevine photosynthesis and stomatal conductance was measured to determine if the tetralone-ABA affected growing season physiology. A Ll-600 Porometer/Fluorometer (LI-COR Biosciences, NE, US) was used to quantify the photosynthetic efficiency, expressed as *ETR* (µmol·m^-2^·s^-1^), calculated from operating efficiency of PSII (Φ*_PSII_*) and ambient light condition (*Q_amb_*), and stomatal conductance, expressed as *g_sw_* (mol·m^-2^·s^-1^). Harvest yield data, including fruit yield per vine and number of clusters per vine, were obtained during the harvest in both years.

### 2.7 RNA-seq during deacclimation assays, library preparation and data processing

During each deacclimation assay conducted in the 2022-2023 experiment and in parallel with bud cold hardiness measurements, bud samples were collected for RNA-seq to investigate the transcriptome effect of tetralone-ABA during deacclimation. Three biological replicates, each composed of five pooled buds, were used for extraction and library preparation. The Cornell University Institute of Genomic Diversity (Ithaca, NY, USA) provided technical support for library construction using Lexogen QuantSeq 3’mRNA-Seq Prep Kit (Lexogen, Greenland, NH). Sequencing of libraries was conducted at Cornell University Institute of Biotechnology (Ithaca, NY) using NextSeq500 (Illumina, Inc., San Diego, CA, USA) with 95 samples per lane. The read length was 85 bp, and sequencing was replicated three times in each library for technical validity.

RNA-seq data was analyzed following a previously generated pipeline specifically optimized for grapevine [54]. Standardized library QC (FastQC), trimming (BBDuk) was conducted for quality control of the libraries, and transcript alignment was conducted in STAR using *V. vinifera* 12X.v2 genome and VCost.v3 annotation [55,56]. The resulting gene count matrix, composed by libraries (as columns) and genes (as rows) was analyzed using DESeq2 to identify differentially expressed genes [57]. Principal Component Analysis (PCA), Uniform Manifold approximation and Projection (UMAP), and Weighted Gene Co-expression Network Analysis (WGCNA) were used for the characterization of transcriptome and the identification of gene co-expression modules [58]. After the identification of target gene co-expression modules that showed significant correlation with deacclimation and responded to tetralone-ABA treatment, we applied a two-step filtering to identify target genes. As the first step, the genes exhibiting a module membership (Pearson correlation coefficient of gene expression with the target module eigengene (ME), the PC1 of all the genes in a module) of greater than 0.8 were reserved. This step ensured that the selected genes follow the pattern of target gene co-expression modules. As the second step, we retained genes with a false discovery rate (FDR) ≤ 1e-30 in the differential expression analysis using DESeq2, with the ME values of target modules as the only factors in the gene expression model. This step ensured a highly significant correlation between gene expression and the ME values of target modules. The resulting genes closely followed the behavior of the ME values of the target gene co-expression modules, making them suitable for downstream analysis. Enriched pathways among the target genes were detected through over representation analysis (ORA) using VitisNet function pathways [59,60]. Significantly enriched pathways were examined for correlations with known biological functions, and hub genes were determined based on the synergy of statistical significance and biological functionality. In addition, we also manually examined individual genes, or the genes involved in pathways that were previously reported to be functional in grapevine cold hardiness.

While the primary analysis incorporated all libraries, additional analyses were conducted on libraries from each deacclimation assay to identify tetralone-ABA-induced differential expression. These analyses used ‘days in growth permissive condition’ and ‘treatment’ as factors in the gene expression model to capture specific responses in early, mid-, and late-winter buds.

## 3. Results

### 3.1 Dormant season physiology

The impact of tetralone-ABA treatment on the relative chlorophyll concentration (SPAD), dormant season cold hardiness and deacclimation rate of three cultivars is shown in Figure 1. For ‘Riesling’, tetralone-ABA treated vines started showing significantly lower SPAD one-day post-treatment (10/06/2022) (Figure 1A). The treatment effect was also observed in the following three measurements at one-week post-treatment (10/12/2022), two-week post-treatment (10/19/2022) and three-week post-treatment (10/26/2022). Visually, tetralone-ABA treated ‘Riesling’ vines showed a yellower pattern on one-week and two-week post-treatment and reached 100% defoliation at three-week post-treatment, as the control vines just started defoliation (Figure 1A). In ‘Cayuga White’, tetralone-ABA treatment effect was limited to a marginally lower SPAD at only one-week post-treatment (Figure 1A). Defoliation occurred simultaneously in tetralone-ABA treated and control ‘Cayuga White’ vines. In ‘Aravelle’, the tetralone-ABA treatment effect was intermediate to its parents: significant reductions in SPAD were noted at one-week and two-week post-treatment, yet defoliation timings were equivalent in tetralone-ABA treated and control vines (Figure 1A).

**Figure 1.**
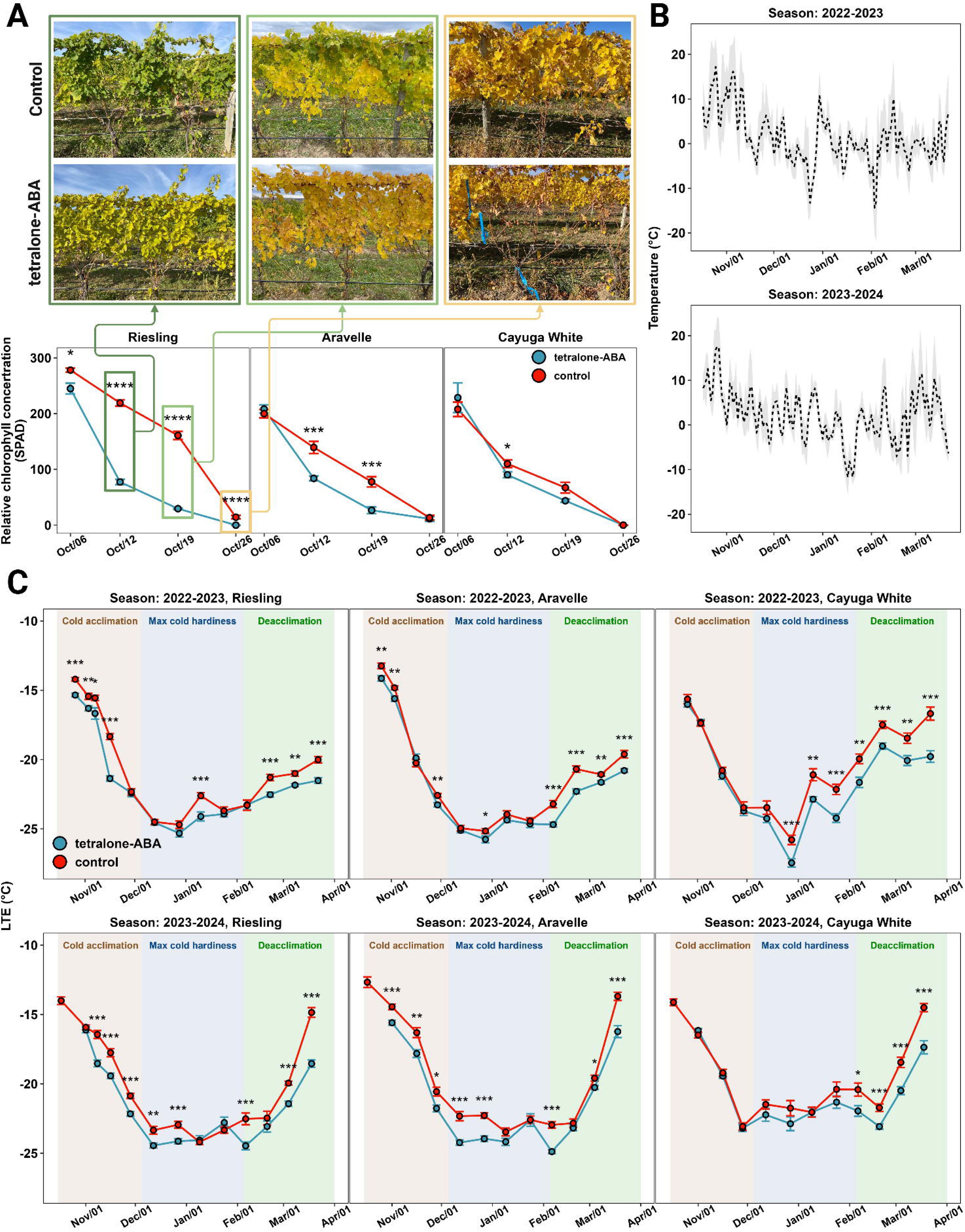
The impact of tetralone-ABA on grapevine relative chlorophyll concentration and dormant season physiology. A) Relative chlorophyll concentration of ‘Riesling’ after treatment application on 10/05/2022. Images represent changes in ‘Riesling’ vines during each relative chlorophyll concentration measurement. B) Daily temperature in the 2022-2023 and 2023-2024 dormant seasons, with dashed lines representing mean temperatures and shaded areas indicating maximum and minimum temperature ranges. C) Bud cold hardiness of three cultivars in the 2022-2023 and 2023-2024 dormant seasons. The mean of bud LTE is shown with the error bars representing standard error (n ≥ 15).

While the air temperature showed a general ‘U’-shaped pattern during the two dormant seasons, the 2023-2024 dormant season was on average, warmer and exhibited less temperature fluctuation as compared to the 2022-2023 dormant season (Figure 1B). As a result, the chilling accumulation occurred much faster under field conditions during the 2023-2024 dormant season compared with 2022-2023 (Figure S1). Regarding cold hardiness, a notable difference between the two seasons is that the deacclimation process occurred much faster (steeper loss of cold hardiness) in the 2023-2024 dormant season regardless of cultivar or treatment (Figure 1C). For ‘Riesling’ and in both dormant seasons, significant enhancement in bud cold hardiness were noted in tetralone-ABA treated vines during cold acclimation following decreasing minimum temperature from mid-October to mid-November (Figure 1C). Deacclimation was observed in control vines following a warming event in early January 2023, and tetralone-ABA treated vines exhibited superior bud cold hardiness in this period (Figure 1C). In both dormant seasons, significant enhancement of bud cold hardiness was observed for tetralone-ABA treated vines after mid-February, demonstrating tetralone-ABA’s delaying effect on field deacclimation (Figure 1C). To compare, in ‘Cayuga White’, no treatment effect was identified during cold acclimation in two seasons, but significant enhancements of bud cold hardiness were observed throughout the deacclimation processes until the end of data collection (Figure 1C). The tetralone-ABA treatment effect on ‘Aravelle’ was intermediate to its parents: enhancement of bud cold hardiness was observed in cold acclimation and deacclimation, but the degree of enhancement was comparatively modest (Figure 1C).

In the deacclimation assays conducted under growth permissive conditions, tetralone-ABA showed delayed bud deacclimation responses, and the delay increased with higher field chilling (Figure 2). The control buds collected from early winter exhibited minimal deacclimation rate, measured as 0.2-0.3 °C·day^-1^ (Figure 2). Tetralone-ABA treatment did not alter the overall deacclimation rate of these early winter collections (Figure 2). The control buds collected from mid-winter and late winter showed increased deacclimation rate, while tetralone-ABA treatment consistently delayed the loss of cold hardiness at later timepoints in these assays, resulting in a decreased deacclimation rate (Figure 2). On average, the deacclimation rate of mid and late winter deacclimation assays was decreased by tetralone-ABA treatment by 25% relative to control.

**Figure 2.**
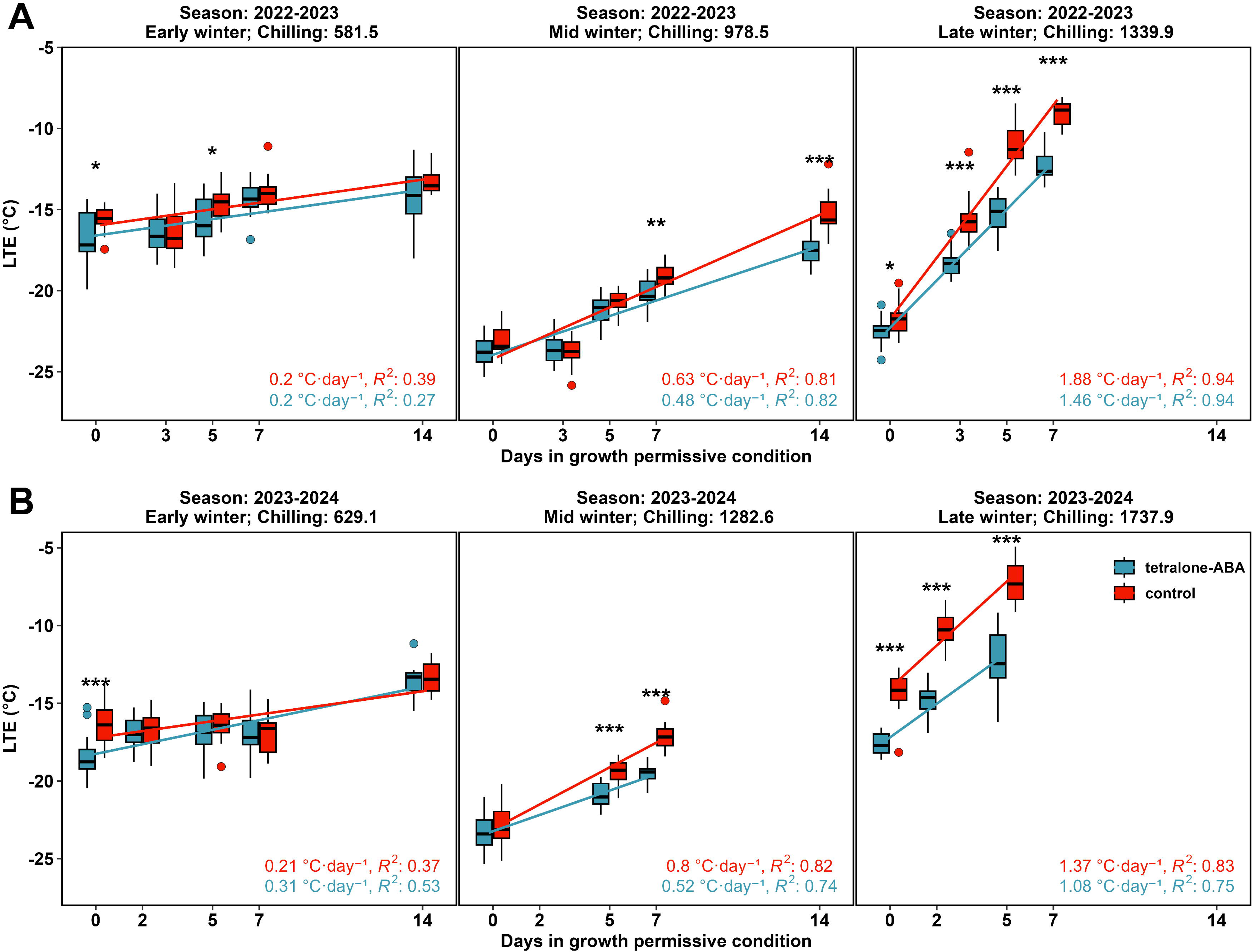
The impact of tetralone-ABA on the deacclimation of ‘Riesling’ buds in controlled environment. A) Deacclimation assays conducted in the 2022-2023 dormant season. B) Deacclimation assays conducted in the 2023-2024 dormant season.

### 3.2 Transcriptomic analysis of deacclimation and tetralone-ABA treatment impact

In parallel with each deacclimation assay conducted in the 2022-2023 dormant season, replicate buds were examined for their transcriptomic response to tetralone-ABA during deacclimation. A total of 90 libraries were sequenced from the three deacclimation assays. Each library met FastQC per base quality standards. On average, each library contained 4.6 million reads, with a unique mapping rate averaging 78.9%. Six libraries were identified as outliers in PCA, and these libraries were removed for downstream analysis (Figure S2). After filtering out low count genes (genes with less than five transcripts per library), the final gene count matrix for any downstream analysis was composed of 84 libraries and 15,002 genes (35% of total transcriptome).

We applied PCA, UMAP and WGCNA to the final gene expression matrix for the overall transcriptome characterization of the libraries. The differentiation of the libraries from different treatment, timepoints and deacclimation assays using PC1 and PC2 from PCA and UMAP1 and UMAP2 from UMAP is shown in Figure S3. The visualization of PC1 and UMAP2 indicates that the libraries generated from early and mid-winter buds did not show significant changes during the deacclimation assays, and there was limited treatment effect of tetralone-ABA (Figure 3A and B). The analysis of the first deacclimation assay libraries resulted in no differentially expressed genes between the two treatments. The analysis on the second deacclimation libraries assay resulted in 133 genes that exhibited differential expression at FDR ≤ 0.01 between the two treatments. Among the 108 annotated genes in these genes, 73 genes exhibited higher expression in tetralone-ABA treated buds, 35 genes exhibited higher expression in control buds (Figure S4). No pathway was statistically enriched among these genes. We applied WGCNA on these 108 genes and identified three major expression patterns during the second deacclimation assay: 1) increase and converge (50 genes in module ‘turquoise’), in which gene expression was initially higher in ABA-treated buds and was later increased during the experiment, but the increase was slower compared to control, leading to the convergence of gene expression of the two treatments by the end of the experiment; 2) decrease and converge (23 genes in module ‘brown’), in which the above pattern was reversed; 3) two directions and converge (35 genes in module ‘blue’), in which gene expression was initially higher in control buds but was decreased during the experiment when the expression was increased in tetralone-ABA treated buds, which led to the convergence of gene expression of the two treatments by the end of the experiment (Figure S5).

**Figure 3.**
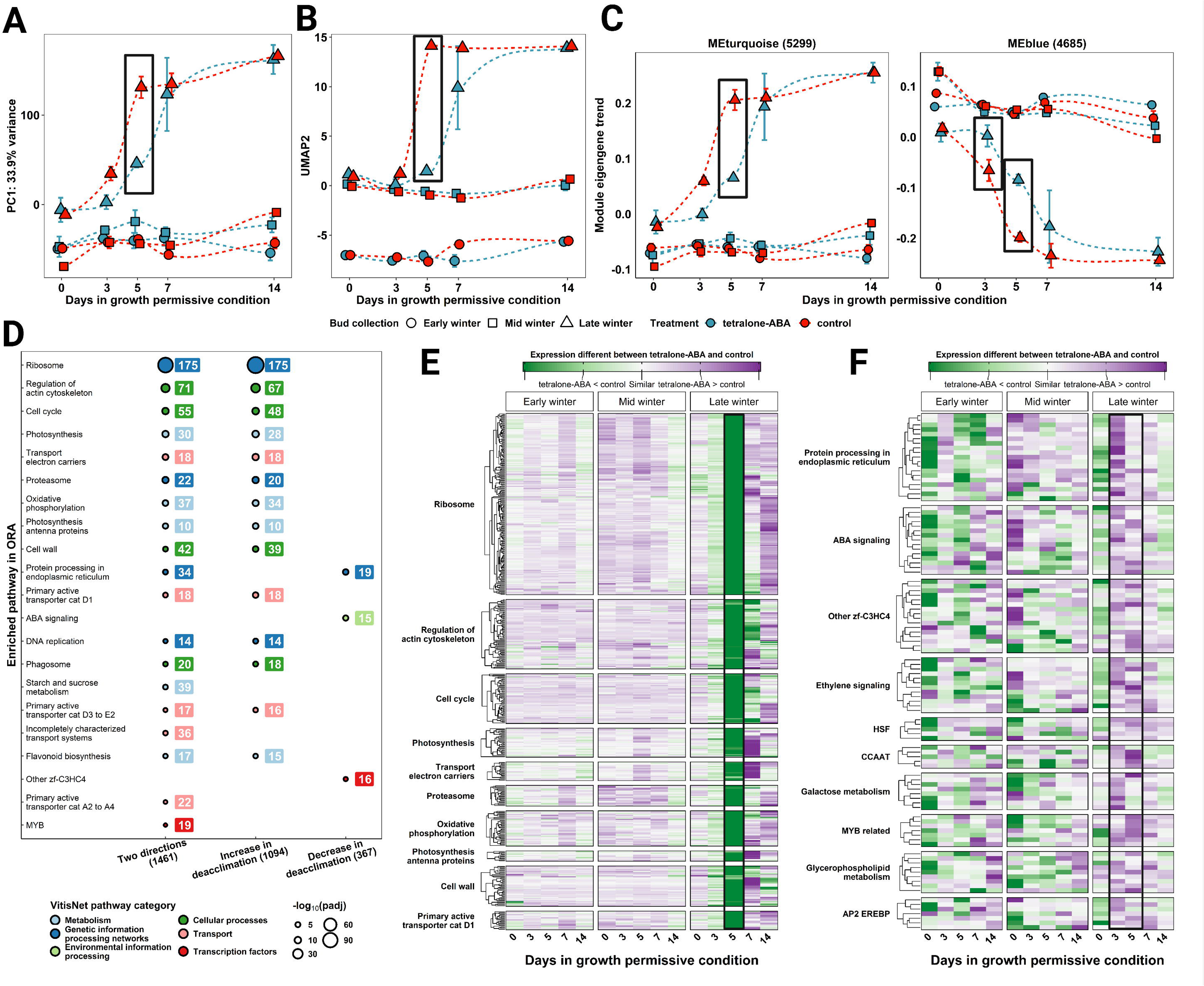
Identification of two gene clusters that correlated with deacclimation and responded to tetralone-ABA treatment. A) Behavior of PC1 from the PCA of the active transcriptome in the three deacclimation assays. B) Behavior of UMAP2 from the UMAP of the active transcriptome in the three deacclimation assays. C) Expression pattern of the module eigengenes (MEs) in modules ‘turquoise’ and ‘blue’ that correlated with deacclimation and responded to tetralone-ABA treatment. D) Pathway enrichment analysis of the genes showed significant correlation with deacclimation and responded to tetralone-ABA treatment, categorized by VitisNet pathway system. E) Average expression difference between control and tetralone-ABA of the ‘increase in deacclimation’ genes. The greatest expression differences (highlighted) were observed on day 5 during the late winter deacclimation assay. F) Average expression difference between control and tetralone-ABA of the ‘decrease in deacclimation’ genes.

In contrast, the PC1 and UMAP2 of the libraries generated from late winter buds showed a rapid increase between day 3 and day 5 and generally plateaued at a higher value after 7 d (Figure 3A and B). The same increase was observed in the sample collected from tetralone-ABA treated buds, but the increase was delayed (Figure 3A and B). The WGCNA using all the genes in the final gene count matrix identified 11 gene co-expression modules, excluding module ‘grey’ containing noisy genes. The number of genes in each module ranged from 57 in module ‘greenyellow’ to 5299 in module ‘turquoise’. The MEs of all the gene co-expression modules are depicted in Figure S6. Notably, the MEs of the two largest modules, ‘turquoise’ (5299 genes) and ‘blue’ (4685 genes) displayed divergent behaviors during the experiment, and their behaviors are either nearly identical or opposite to PC1 from PCA (Figure 3A and C). These genes represent nearly 2/3 of the active transcriptome identified in our experiment. In summary, there were significant differences of the transcriptomes of the buds under tetralone-ABA treatment and control, especially on day 3 and day 5 in the third deacclimation assay when the buds were rapidly deacclimating. Two gene clusters have expression patterns closely linked to deacclimation and are impacted by tetralone-ABA treatment.

### 3.3 Pathways impacted in deacclimation and in response to tetralone-ABA treatment in late winter buds

After gene filtering, we identified 1461 genes of interest associated with deacclimation from gene co-expression modules ‘turquoise’ and ‘blue’. Among these genes, 1094 genes showed increasing expression during deacclimation (same trend as ME ‘turquoise’), and 367 genes showed decreasing expression (same trend as ME ‘blue’). The statistics of the differential expression analysis and detailed information of these genes are available in Data S1 and S2. Enrichment analysis identified 20 pathways in VitisNet that were significantly enriched at FDR ≤ 0.001 (Figure 3D). Genes which significantly increased during deacclimation were associated with pathways include several pathways in genetic information processing networks, cellular processes, transport and metabolism (Figure 3D). The most enriched pathways, including ribosome (vv23010), regulation of actin cytoskeleton (vv44810), cell cycle (vv44110), photosynthesis (vv10195), transport electron carriers (vv50105), oxidative phosphorylation (vv10190) and cell wall (vv40006) are essential pathways for plant growth. The genes in these pathways showed limited expression change and differences between two treatments in the first and second deacclimation assays with early winter and mid-winter buds, respectively (Figure 3E). However, in the third deacclimation assay with late winter buds, significantly lower expression was identified in tetralone-ABA treatment on day 3 and day 5 in almost every gene in every enriched pathway (Figure 3E). Genes which significantly decreased during deacclimation were associated with pathways include protein processing in endoplasmic reticulum (vv24141), ABA signaling (vv30001), ethylene signaling (vv30008), HSF (heat shock factor, vv60037) and galactose metabolism (vv10052). The genes in these pathways did not show consistent differential expression in the first and second deacclimation assays but showed consistent higher expression on day 3 and day 5 in tetralone-ABA treated buds in the third deacclimation assay (Figure 3F).

### 3.4 Detailed investigation of enriched and functional pathways

In addition to these analyses, we investigated some enriched pathways in detail and manually evaluated some other pathways that were reported to be functional in grapevine cold hardiness and deacclimation.

#### ABA biosynthesis and signaling pathways

ABA signaling pathway was one of the significantly enriched pathways among the genes that showed decreased expression during deacclimation (Figure 3F). However, most of the genes with observable expressions in ABA metabolism in VitisNet did not reveal notable treatment effects (Figure S7). Among the seven copies of the gene encoding for 9-cis-epoxycarotenoid dioxygenase (NCED, key enzyme for ABA biosynthesis), three genes passed low count filtering in our experiment. None of the three genes revealed any treatment or chilling effect (Figure S7). Among the four copies of the gene encoding for ABA 8’-hydroxylase (key enzyme for ABA degradation), two genes passed low count filtering in our experiment. One gene (*VIT_03s0063g00380*) showed slightly higher expression in tetralone-ABA treated late winter buds under field condition (day 0), but no difference under growth permissive condition (day 3 to day 14) (Figure S8A). Both copies of the gene encoding for ABA glucosidase (key enzyme for ABA conjugation) passed low count filtering in our experiment. One gene (*VIT_17s0000g02680*) showed generally increasing expression with deacclimation, and the increase was delayed by tetralone-ABA on day 5 in the third deacclimation assay with late winter buds (Figure S8B).

In ABA signaling, a group of genes encoding key components of the signaling and ABA responsive genes not only showed decreasing expression with deacclimation but also revealed the tetralone-ABA treatment effect in delaying this decrease, especially in the third deacclimation assay with late winter buds (Figure S8C). These genes include one encoding for AREB2, two for RCAR, one for PP2C and two for SnRK2 in ABA signaling, along with six in ABA responsive genes, including a highly expressed member of *EIN3* (Figure S8C).

#### Sugar metabolism pathways

A total of 16 genes in galactose metabolism pathway were identified for their expression correlation with deacclimation and tetralone-ABA. Among these genes, three genes encoding for beta-galactosidase were upregulated in the third deacclimation assay with late winter buds, and the upregulation was delayed by tetralone-ABA treatment (Figure S9). Four genes encoding for galactinol synthase were downregulated in the third deacclimation assay, and the downregulation was delayed by tetralone-ABA (Figure S9). A total of 36 genes in starch and sucrose metabolism pathways (after excluding the overlapping genes in galactose metabolism pathway) were identified for their expression correlation with deacclimation and tetralone-ABA (Figure S10). Among these genes, 13 genes associated with the proteins related to endo-1,3-beta-glucosidase and one gene encoding for callose synthase catalytic subunit were upregulated in the third deacclimation assay with late winter bud, and the upregulation was delayed by tetralone-ABA in most of these genes (Figure S10).

#### Cell wall pathway

In the cell wall pathway, the expression of 42 genes correlated with deacclimation and showed response to tetralone-ABA. Of these, 39 genes were upregulated in the third deacclimation assay, and upregulation was delayed by tetralone-ABA (Figure S11). These proteins can be categorized into two functional groups: cell wall proteins and cell wall substrate modification enzymes. The cell wall protein group includes unannotated hydroxyproline-rich glycoproteins (six genes), fasciclin-like arabinogalactan-protein (FLA, three genes) and leucine-rich repeat protein/extensin (LRX, four genes) (Figure S11). The cell wall substrate modification enzyme group includes endo-1,4-beta-glucosidase (also known as cellulase, five genes), pectate lyase (PL, two genes), pectin-acetylesterase (PAE, six genes), polygalacturonase (PG, also known as pectinase, seven genes) and xyloglucan endotransglucosylase/hydrolase (XTH, two genes) (Figure S11).

#### ICE, CBF and DREB

Among all the 14 *Inducers of CBF Expression* (*ICE*) and *CBF/DREB* genes annotated in VitisNet, five genes passed the low count filtering, and none of these genes are *CBF* (Figure S12). One gene encoding DREB (*VIT_13s0067g01960*) was downregulated during deacclimation, and tetralone-ABA delayed the downregulation in the third deacclimation assay with late winter buds (Figure S12). One gene encoding ICE (*VIT_14s0068g01200*) was upregulated during deacclimation, and tetralone-ABA delayed the upregulation in the third deacclimation assay (Figure S12).

### 3.5 Budbreak phenology

In the 2022-2023 experiment, field phenology recording began on 04/19/2023. A significant delay in phenology was observed in tetralone-ABA treated vines for all three cultivars, starting from the initial data collection and continuing until late May when clear inflorescences developed (Figure 4A). The dates of budbreak, defined as when more than 50% of experimental vines reached the budbreak stage (Modified Shaulis Field Score 4), were 04/24/2023, for ‘Riesling’ and ‘Aravelle,’ and 05/01/2023, for ‘Cayuga White’ in the control vines (Figure 4A). In the tetralone-ABA treated vines, budbreak occurred on 05/05/2023, for ‘Riesling,’ 05/03/2023, for ‘Aravelle’, and 05/09/2023, for ‘Cayuga White,’ representing delays of 11, 9, and 8 d, respectively, compared to the control vines (Figure 4A). In the 2023-2024 experiment, field phenology recording began on 04/23/2024. Similarly, significant delay in phenology was observed in tetralone-ABA treated vines for all three cultivars until when clear inflorescences developed, but the difference of the numeric phenological stage between two treatments was generally smaller as compared to the previous year (Figure 4A). The dates of budbreak were 05/01/2024, for ‘Riesling’ and ‘Aravelle,’ and 05/03/2024, for ‘Cayuga White’ in the control vines (Figure 4A). In the tetralone-ABA treated vines, budbreak occurred on 05/05/2024, for ‘Riesling and ‘Aravelle,’ and 05/07/2023, for ‘Cayuga White,’ representing a delay of 4 d, compared to the control vines in all three cultivars (Figure 4A).

**Figure 4.**
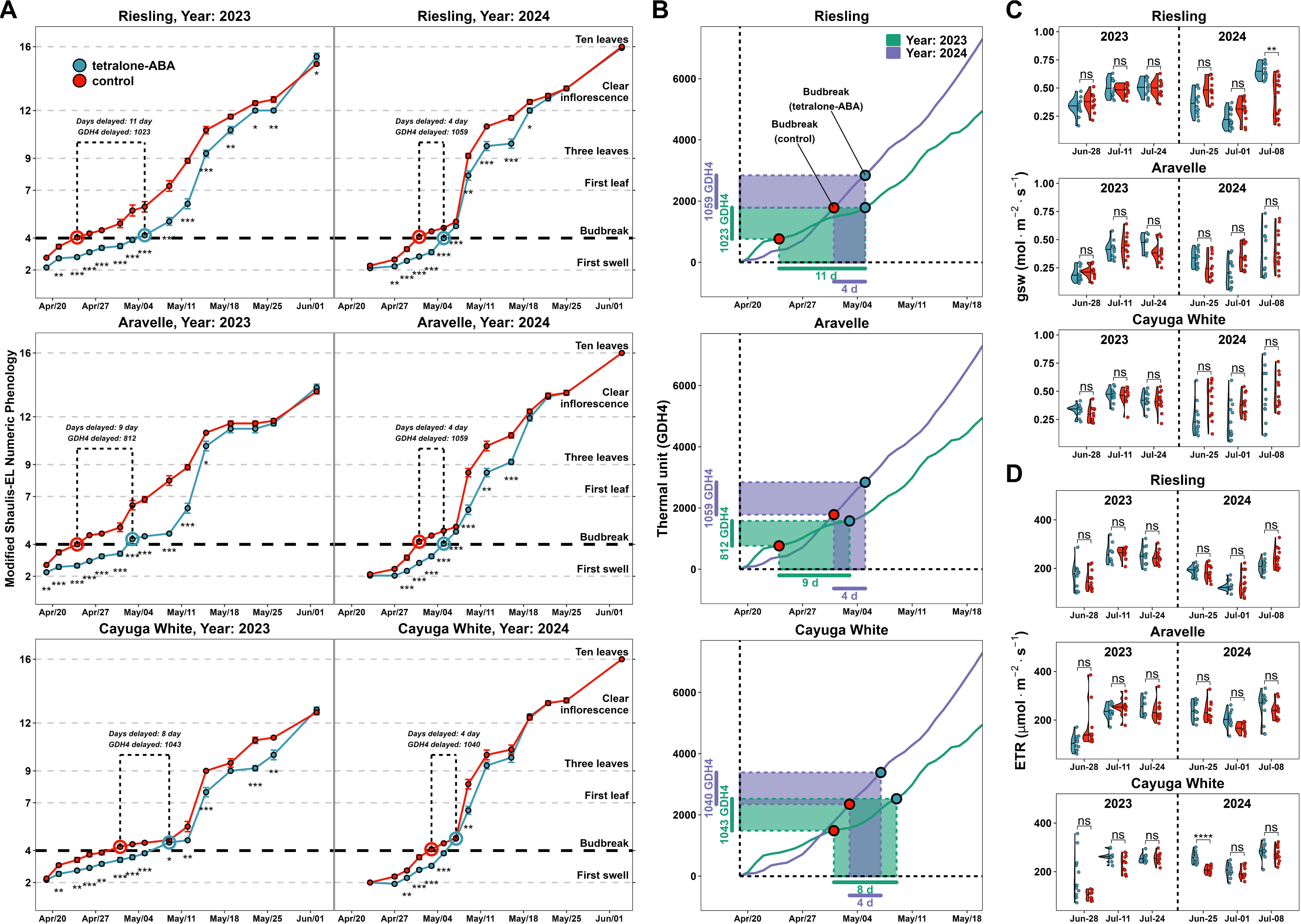
The impact of tetralone-ABA on grapevine phenology and growing season physiology. A). The phenology of the three cultivars during the early growing season in 2023 and 2024. Error bar represents standard error (n = 12). All the observations after the last observation shown in the figure showed no difference between the two treatments. B) Comparison between day and GDH4 (growing degree hours using 4 °C as the base temperature) as a measurement for tetralone-ABA induced delay of budbreak. C) Stomatal conductance expressed as *g_sw_* (mol·m^-2^·s^-1^). D) Relative photosynthesis efficiency expressed as *ETR* (µmol·m^-2^·s^-1^).

While the delay in days exhibited substantial differences between the two years, we observed a significant difference in weather conditions: heat accumulation (calculated in GDH4) occurred much faster in the early spring of 2024 when grapevines were actively deacclimating and initiating growth resumption as compared to 2023 (Figure 4B). When transforming the delay (the gap in budbreak timing between control vines and tetralone-ABA treated vines) from days to GDH4, a more precise measurement of thermal units, the delay became much more consistent between the two years (Figure 4B). The delays from tetralone-ABA treatment ranged narrowly between 812 GDH4 to 1059 GDH4 across cultivars and years (Figure 4B).

### 3.6 Growing season physiology and harvest yield

During the growing season in 2023, we measured photosynthetic efficiency and stomatal conductance on 06/28/2023, 07/11/23 and 07/24/2023. Neither physiological parameter showed a difference between the two treatments in any of the three cultivars (Figure 4C and D). During the growing season in 2024, we measured photosynthetic efficiency and stomatal conductance on 06/25/2024, 07/01/24 and 07/08/2024. Except for the differences between the two treatments in *g_sw_* of ‘Riesling’ on 07/08/2024 and *ETR* of ‘Cayuga White’ on 06/25/2024, neither physiological parameter showed a difference between the two treatments in any of the three cultivars (Figure 4C and D).

In 2023, the experimental vines were harvested on 10/10/2023 for ‘Riesling’ and 09/26/2023 for ‘Aravelle’. In 2024, the experimental vines were harvested on 10/06/2024 for ‘Riesling’ and 09/16/2024 for ‘Aravelle’ and ‘Cayuga White’. Harvest data, including clusters per vine and yield per vine, were collected to evaluate the treatment effect. The average cluster weight per vine was calculated based on clusters per vine and yield per vine. No differences were identified between the two treatments across the cultivars in two years (Figure S13).

## Discussion

With changing climate and shifts in frost risk to perennial fruit crops, it is critical to develop and evaluate potential mitigation methods for reducing deacclimation risk and delaying budbreak phenology. This research represents a detailed exploration of the effects of tetralone-ABA treatment as a novel sprayable product to enhance grapevine resilience in the dormant season of three grapevine cultivars: ‘Riesling’, ‘Aravelle’, and ‘Cayuga White’. By analyzing the nuances of how such a treatment affects grapevine at both the molecular and physiological levels, this study contributes to a more comprehensive understanding of grapevine biology, offering potential insights for enhancing grapevine resilience and informing sustainable viticultural strategies in an era of increasing environmental challenges under climate change.

Many studies have explored different forms of ABA analogs for their feasibility to enhance grapevine bud cold hardiness in dormant season. To date, the most tested ABA analog is 8’-acetylene ABA, which is synthesized by replacing the 8’methyl group in ABA with an acetylene, avoiding this analog being the target of ABA catabolism pathway [61]. Some studies report significant enhancement of grapevine bud cold hardiness from post-harvest application of this analog, yet others reported limited effect [36,39,40]. Tetralone-ABA is another form of ABA analog that is synthesized by introducing an aromatic ring in place of the vinyl methyl group in the endogenous ABA, allowing for slower catabolism thus potentially maintaining prolonged bioactivity [38,62]. Previous studies of tetralone-ABA’s effect on grapevine indicate that this analog, even though synthesized through a different method (ABA-1016), enhanced bud cold hardiness close to the end of the dormant season, suggesting a potential role in slowing deacclimation [39,40].

In our study, we evaluated the efficacy of a new tetralone-ABA product, ABA-1102 in enhancing cold hardiness and delaying spring budbreak of three grapevine cultivars: ‘Riesling’, ‘Cayuga White’, and their progeny, ‘Aravelle’. Shortly after tetralone-ABA application, we observed accelerated chlorophyll degradation and early defoliation in tetralone-ABA treated vines, however, the effect varies between different cultivars (Figure 1A). This result contrasts with previous studies that reported no impact of tetralone-ABA on leaf senescence [39]. Effects were strongest for ‘Riesling’ while ‘Cayuga White’ has a much more limited reduction in SPAD readings (Figure 1A). ‘Aravelle’, the offspring of ‘Riesling’ and ‘Cayuga White’, exhibited intermediate treatment effect compared to its parents (Figure 1A). This cultivar specific response is intriguing as it suggests there may be differences in sensitivity to tetralone ABA that differs between the *V. vinifera* and *V. hybrid* genetic backgrounds. The intermediate phenotype in ‘Aravelle’ supports this hypothesis, though many more cultivars and pedigrees would need to be evaluated to fully explore this response.

For two dormant seasons, tetralone-ABA significantly enhanced the bud cold hardiness of ‘Riesling’ during early winter in the process of initial cold acclimation but had no effect on ‘Cayuga White’ in the same period (Figure 1C). In contrast, tetralone-ABA significantly enhanced the bud cold hardiness of ‘Cayuga White’ in late winter and during deacclimation, but the enhancement was less intensive in ‘Riesling’ (Figure 1C). Much like the senescence response, tetralone-ABA’s effect on enhancing bud cold hardiness in ‘Aravelle’ was intermediate to its effect on its parents (Figure 1C). The intermediate response to tetralone-ABA in ‘Aravelle’, compared to its parents, potentially indicates that tetralone-ABA’s impact bud cold hardiness is also an inheritable quantitative trait, which is controlled by unknown genetic variations among *Vitis*. High sensitivity to tetralone-ABA may be a preferred trait for future breeding programs as it might facilitate the utilization of different forms of ABA to enhance winter cold hardiness. While enhanced bud cold hardiness during deacclimation as a response to tetralone-ABA has been reported in previous studies [36,39,40], this is the first time that enhanced bud cold hardiness during cold acclimation was observed, especially on ‘Riesling’, an economically important cultivar for the cool climate viticultural regions in North America.

Tetralone-ABA had a clear effect on decreasing deacclimation rate in the deacclimation assays conducted on the buds with more than 1000 chilling units (Figure 2). This result supports observations that tetralone-ABA-induced slower deacclimation observed under field conditions in this study and others [39,40]. Detailed examination of the bud transcriptome during the deacclimation assays indicates diverse transcriptomic regulation differences in control and tetralone-ABA treated vines. No differentially expressed genes were observed in the first deacclimation assay on early winter buds. However, in the second assay on mid-winter buds, 108 genes were differentially expressed between treatments. The differential expression of these genes, especially under field condition, could be the underlying mechanism of the decreased deacclimation rate observed in the second deacclimation assay. Although no pathway was significantly enriched among these genes, we noticed several genes in the pathways that were previously reported to have crucial biological functions in the dormancy or cold hardiness of grapevines. These genes include three genes encoding enzymes/proteins in ABA signaling (*VIT_14s0030g00690* encoding for ABI3, *VIT_15s0046g01050* encoding for RCAR1 and *VIT_13s0067g02130* encoding for ERD15), one gene in ethylene signaling (*VIT_05s0020g04180* encoding for CTR1), two genes in flower development (*VIT_01s0011g02120* encoding for HUA2 and *VIT_02s0025g04390* encoding for SEU1), five genes encoding for ribosome subunits, three highly expressed genes encoding for heat shock proteins, two genes encoding for histone metabolism, nine genes encoding for various transcription factors and six genes encoding for cell membrane transporters or channels (Figure S4).

Two large groups of genes demonstrated expression patterns that correlated with deacclimation and tetralone-ABA in the third deacclimation assay. In control, the expression of these genes remained relatively unchanged in the first and second deacclimation assays conducted on early winter buds and mid-winter buds but showed significant increase/decrease in the third deacclimation assay with late winter buds when buds were rapidly deacclimating (Figure 2 and 3). The observed unchanged gene expression patterns in early and mid-winter buds likely indicates that the grapevines were either under endodormancy or transitioning from endodormancy to ecodormancy, during which buds are resistant to deacclimation or unable to initiate growth even under growth-permissive conditions. In contrast, the significant changes in gene expression observed during the late winter deacclimation assay likely indicates that, after the satisfaction of chilling requirement, the grapevines had fully transitioned to ecodormancy. This period is characterized by an increased ability of buds to respond to external stimuli, such as temperature cues, by deacclimating and preparing for growth [49,63–66]. With tetralone-ABA treatment, the change of gene expression observed in control in the third deacclimation assay, whether and increase or decrease, was significantly delayed (Figure 3). Notably, nearly 2/3 of the active transcriptome of the buds generally followed the above expression pattern and responded to tetralone-ABA treatment. Most of the enriched pathways among the genes that showed increasing expression during deacclimation are fundamental for plant growth, such as ribosome pathway, regulation of actin cytoskeleton pathway, photosynthesis-related pathway and cell cycle pathway (Figure 3D) [67]. The genes with decreasing expression during deacclimation are enriched in the pathways such as protein processing in the endoplasmic reticulum, ABA signaling, ethylene signaling, HSF, and galactose metabolism. ABA signaling, ethylene signaling, and galactose metabolism have been recognized or suggested to play significant roles in the regulation and development of grapevine cold hardiness [32,37,51,54,68,69]. In the protein processing in the endoplasmic reticulum pathway, 17 out of 19 genes encode heat shock proteins. Coupled with the enrichment of HSF, these findings suggest the involvement of heat shock protein biosynthesis downregulation in temperature-regulated deacclimation, aligning with previous findings [51,70]. Regarding the ABA-related pathways, most of the genes involved in ABA metabolism did not show significant response to tetralone-ABA (Figure S7). The only two genes that exhibited significant changes in expression in response to tetralone-ABA are *VIT_03s0063g00380*, encoding for ABA 8’-hydroxylase that degrades ABA, and *VIT_17s0000g02680*, encoding for ABA glucosidase that transforms ABA to inactive form [26]. The higher expression of *VIT_03s0063g00380* in the tetralone-ABA treated buds under field condition may suggest that tetralone-ABA treatment stimulates a response to degrade excessive active ABA-like metabolites as seen in the overexpression lines (Figure S8A) [71]. This finding supports, although indirectly, that tetralone-ABA persists throughout the dormant season in grapevine and at the gene expression level performs as if ABA is being continuously applied to the bud tissue. This explanation also agrees with a previous study that reported detectable tetralone-ABA in grapevine buds by mid-April after the foliar application in the previous fall [40]. Assuming unchanged translation efficiency, the delayed upregulation of *VIT_17s0000g02680* in tetralone-ABA treated buds in the third deacclimation assay suggests the treatment may lead to a higher content of free ABA (Figure S8B). Downstream of ABA synthesis in the genes associated with ABA signaling, many genes encoding for the key proteins for ABA reception and signal transduction were found to be decreasing with deacclimation and the decreasing trend was delayed by tetralone-ABA (Figure S8C). This result indicates that, without tetralone-ABA, grapevine buds tend to downregulate ABA signaling pathway during deacclimation to facilitate the activation of growth-relative pathways, which aligns with previous findings [51,54,72]. With tetralone-ABA, the downregulation of ABA signaling was delayed, thus likely promoting the expression of ABA responsive genes that contribute to prolonged dormancy.

We also manually evaluated several pathways or genes that were previously reported to be important for grapevine dormant season physiology to further evaluate the impact of tetralone-ABA on grapevine transcriptome. These include galactose metabolism pathway, starch and sucrose metabolism pathway, cell wall pathways and ICE, CBF and DREB genes. The *ICE-CBF/DREB-COR* (*COLD REGULATED*) cascade for the genetic control of plant cold stress response is activated with the upregulation of *ICE*, followed by the *ICE*-induced upregulation of *CBF/DREB* that stimulates the expression of *COR*s that encodes functional metabolites to improve cold hardiness [73]. The upregulation of this cascade was repeatedly reported to function in the enhancement of grapevine cold hardiness, however, in none-overwintering tissues such as shoot, leaves and petioles [74–78]. In this study, none of the four copies of *CBF* in *V. vinifera* 12X.v2 genome showed detectable expression in control or tetralone-ABA treated buds in our study. This result disagrees with a previous study that reported thousands fold of increase in the expression of *CBF*s after ABA treatment [25]. Among the three *ICE*s and two *DREB*s with detectable expression, two genes (*VIT_13s0067g01960* and *VIT_14s0068g01200*) revealed significant but divergent expression correlation with deacclimation and tetralone-ABA (Figure S12). The undetectable expression of *CBFs* and the opposite expression patterns of *ICE* and *DREB* potential indicate that these previous findings and conclusions may not be generalizable in dormant grapevine buds [79]. Further targeted gene expression studies need to be done to detail the behavior of *ICE-CBF-COR* cascade in endodormant grapevine tissues.

In the sugar metabolism pathways and the cell wall pathway, we noticed that many genes that showed delayed upregulation under tetralone-ABA only encode for a short list of proteins. These proteins include beta-galactosidase, endo-1,3-beta-glucosidase, various cell wall proteins including hydroxyproline-rich glycoproteins, FLA and LRX, and cell wall substrate modification enzymes including endo-1,4-beta-glucosidase, PL, PAE, PG and XTH (Figure S9, S10 and S11). All these proteins are involved in plant cell wall modification, and their detailed functions and involvement in plant dormancy release are briefed in Note S1. Taking our findings and previous results together, it seems that cell wall remodification is a key process that occurs during deacclimation and dormancy release. Assuming the observed alteration in transcriptome reflects changes in metabolome, we hypothesize a theoretical model of the remodification of cell wall during deacclimation and tetralone-ABA’s impact during this process (Figure S14). During dormancy, plasmodesmata are plugged with callose, which eliminates signal and substrate transduction through the symplastic pathway [80,81]. Cell wall is primarily composed of excessive cellulose, pectin and hemicelluloses and lack cell wall proteins. Such cell walls are structurally stiff but biologically inactive. Upon receiving enough chilling and under growth permissive conditions (e.g., in the third deacclimation assay), endo-1,3-beta-glucosidase (13 genes) is synthesized to degrade the callose in plasmodesma, allowing the restoration of the symplastic pathway. Another group of enzymes, including beta-galactosidases, endo-1,4-beta-glucosidase, PL, PAE, PG and XTH that degrade primary cell wall components are enriched, potentially leading to the loosening of cell wall. In addition, multiple cell wall proteins, including unannotated hydroxyproline-rich glycoproteins, FLA and LRX, are synthesized, which facilitates the signal transduction on cell wall. The cell walls after the above modification are structurally loose and biologically active, thus promoting symplastic substrate transduction and growth restoration (Figure S14). Cell wall loosening has been proposed to be a critical factor in the dormancy release of woody perennials [82,83]. The application of hydrogen cyanamide induces dormancy release on dormant grapevine buds through reactive oxygen species (ROS) and reactive nitrogen species (RNS)-mediated cell wall autodegradation [34,84–86]. With tetralone-ABA, the synthesis of the cell proteins and cell wall substrate degradation enzymes is delayed, which likely leads to delayed loosening and activation of cell wall and, therefore, delayed budbreak.

Under field conditions, we observed a significant delay in budbreak on tetralone-ABA treated vines across three experimental cultivars over two years. When quantified in days, tetralone-ABA delayed budbreak of ‘Riesling’, ‘Aravelle’, and ‘Cayuga White’ by 11, 9, and 8 days respectively in 2023, and by 4 days for all three cultivars in 2024 (Figure 4A). Using GDH4, a more precise measurement of thermal units, the delay remained relatively consistent at approximately 1000 GDH4 across cultivars and years (Figure 4B). These results suggest three key points: 1) nearly six months after tetralone-ABA application on the foliar, there was still sufficient amount of active tetralone-ABA in the vines that could induce significant and consistent delay in budbreak; 2) the delay is likely associated with the resistance to growth when exposed to accumulated heat, and the degree of resistance is likely consistent across cultivars and years; 3) for future studies, GDH4 or other measurements of thermal units or chilling units are better metrics for quantifying the delay or advancement of phenology resulting from treatment. After budbreak, the delaying effect on grapevine growth varied between different cultivars, but no phenological difference could be observed after clear inflorescence is developed. Physiological measurements during the growing season and harvest data showed minimum difference between tetralone-ABA treated vines and control vines (Figure 4C and D). Although we did not conduct fruit quality and wine chemistry analysis, our results indicate that tetralone-ABA treatment effectively induces budbreak delay without negatively impacting growing season physiology and yield (Figure S13).

In conclusion, post-harvest foliar application of tetralone-ABA on grapevines induced faster leaf chlorophyll degradation, earlier defoliation, enhanced cold acclimation, slower deacclimation, and delayed budbreak without affecting growing season physiology or harvest yield. Transcriptome analysis of grapevine buds during deacclimation indicates that multiple fundamental pathways for growth restoration are upregulated during deacclimation, and tetralone-ABA treatment delayed this upregulation. These results align with our hypotheses. The slower deacclimation in tetralone-ABA treated buds observed in both field and controlled environment conditions is likely a consequence of tetralone-ABA’s delaying effect on the regulation of key functional pathways during deacclimation. The results presented in this study not only indicate the feasibility of using tetralone-ABA as a sprayable product to enhance viticultural resilience in dormant season but also contribute to the understanding of the genetic control of grapevine deacclimation. Optimization of treatment application, including rate, timing, and frequency, remains to be conducted. Additionally, since ABA has similar functions in other plant species, further testing of tetralone-ABA’s effectiveness in other perennial fruit crops could help develop novel methods to combat climate change.

## Supporting information

Supporting Material 1 will be used for the link to the file on the preprint site

Supporting Material 2 will be used for the link to the file on the preprint site

## Acknowledgement

The project was primarily funded through 2022 USDA Northeast Sustainable Agriculture Research and Education (SARE) graduate student research grant (GNE22-304), Cornell AgriTech Summer Scholar Program and Cornell University School of Integrative Plant Science, Horticulture Section.

## Data availability

All RNA-seq raw data along with processed gene count matrix and sample metadata are available in NCBI-GEO (accession: GSE273240).

## Author contribution

HW and JPL designed the experiment and conducted the treatment application. HW and JPL collected dormant season physiology data and the RNA-seq data. HW and YP collected phenology and growing season physiology data. YP did RNA extraction and purification. HW and YP analyzed the RNA-seq data. HW and JPL wrote most of the manuscript with contributions from all co-authors.

## References

1. Luedeling E. Climate change impacts on winter chill for temperate fruit and nut production: A review. Sci Hortic 2012;144:218–29.

2. AghaKouchak A, Chiang F, Huning LS et al. Climate Extremes and Compound Hazards in a Warming World. Annu Rev Earth Planet Sci 2020;48:519–48.

3. De Rosa V, Vizzotto G, Falchi R. Cold Hardiness Dynamics and Spring Phenology: Climate-Driven Changes and New Molecular Insights Into Grapevine Adaptive Potential. Front Plant Sci 2021;12:591.

4. Gunathilaka RPD, Smart JCR, Fleming CM. The impact of changing climate on perennial crops: the case of tea production in Sri Lanka. Clim Change 2017;140:577–92.

5. Evans RG. The art of protecting grapevines from low temperature injury. Proceedings Of ASEV 50th Anniversary Annual Meeting. Seattle WA, 2000, 60–72.

6. Zabadal TJ, Dami IE, Goffinet MC et al. Winter Injury to Grapevines and Methods of Protection. Michigan State University Extension, 2007.

7. Poling EB. Spring cold injury to winegrapes and protection strategies and methods. HortScience 2008;43:1652–62.

8. Warmund MR, Guinan P, Fernandez G. Temperatures and Cold Damage to Small Fruit Crops Across the Eastern United States Associated with the April 2007 Freeze. HortScience 2008;43:1643–7.

9. Londo JP, Moyer MM, Mireles M et al. Evaluation of Sample Preparation Practices Common with Differential Thermal Analysis of Grapevine Bud Cold Hardiness. Am J Enol Vitic 2023;74:0740002.

10. Martinson TE, White GB. Estimate of crop and wine losses due to winter injury in the finger lakes. 2004.

11. Dami I, Lewis D. 2014 Grape Winter Damage Survey Report. Ohio Agricultural Research and Development Center, 2014.

12. Poni S, Sabbatini P, Palliotti A. Facing Spring Frost Damage in Grapevine: Recent Developments and the Role of Delayed Winter Pruning – A Review. Am J Enol Vitic 2022;73:211–26.

13. Wang H, Dami IE. Evaluation of Budbreak-Delaying Products to Avoid Spring Frost Injury in Grapevines. Am J Enol Vitic 2020;71:181–90.

14. Fuller MP, Hamed F, Wisniewski M et al. Protection of plants from frost using hydrophobic particle film and acrylic polymer. Ann Appl Biol 2003;143:93–8.

15. Dami IE, Beam BA. Response of Grapevines to Soybean Oil Application. Am J Enol Vitic 2004;55:269–75.

16. Centinari M, Gardner DM, Smith DE et al. Impact of Amigo Oil and KDL on Grapevine Postbudburst Freeze Damage, Yield Components, and Fruit and Wine Composition. Am J Enol Vitic 2018;69:77–88.

17. Loseke BA, Read PE, Blankenship EE. Preventing spring freeze injury on grapevines using multiple applications of Amigo Oil and naphthaleneacetic acid. Sci Hortic 2015;193:294–300.

18. Persico MJ, Smith DE, Centinari M. Delaying Budbreak to Reduce Freeze Damage: Seasonal Vine Performance and Wine Composition in Two *Vitis vinifera* Cultivars. Am J Enol Vitic 2021:ajev.2021.20076.

19. Reisch BI, Owens CL, Cousins PS. Grape. In: Badenes ML, Byrne DH (eds.). Fruit Breeding. Boston, MA: Springer US, 2012, 225–62.

20. Gutierrez B, Schwaninger H, Meakem V et al. Phenological diversity in wild and hybrid grapes (Vitis) from the USDA-ARS cold-hardy grape collection. Sci Rep 2021;11:24292.

21. Zhang Y, Dami IE. Foliar Application of Abscisic Acid Increases Freezing Tolerance of Field-Grown Vitis vinifera Cabernet franc Grapevines. Am J Enol Vitic 2012;63:377–84.

22. Li S, Dami IE. Responses of Vitis vinifera ‘Pinot gris’ Grapevines to Exogenous Abscisic Acid (ABA): I. Yield, Fruit Quality, Dormancy, and Freezing Tolerance. J Plant Growth Regul 2016;35:245–55.

23. Sun X, Zhao T, Gan S et al. Ethylene positively regulates cold tolerance in grapevine by modulating the expression of ETHYLENE RESPONSE FACTOR 057. Sci Rep 2016;6:24066.

24. Vergara R, Noriega X, Aravena K et al. ABA Represses the Expression of Cell Cycle Genes and May Modulate the Development of Endodormancy in Grapevine Buds. Front Plant Sci 2017;8:812.

25. Rubio S, Noriega X, Pérez FJ. Abscisic acid (ABA) and low temperatures synergistically increase the expression of CBF/DREB1 transcription factors and cold-hardiness in grapevine dormant buds. Ann Bot 2019;123:681–9.

26. Abrams SR, Loewen MC. Chemistry and chemical biology of ABA. Advances in Botanical Research. Vol 92. Elsevier, 2019, 315–39.

27. Basu S, Rabara R. Abscisic acid — An enigma in the abiotic stress tolerance of crop plants. Plant Gene 2017;11:90–8.

28. Liu J, Sherif SM. Hormonal Orchestration of Bud Dormancy Cycle in Deciduous Woody Perennials. Front Plant Sci 2019;10, DOI: 10.3389/fpls.2019.01136.

29. Yang Q, Gao Y, Wu X et al. Bud endodormancy in deciduous fruit trees: advances and prospects. Hortic Res 2021;8:139.

30. Zheng C, Halaly T, Acheampong AK et al. Abscisic acid (ABA) regulates grape bud dormancy, and dormancy release stimuli may act through modification of ABA metabolism. J Exp Bot 2015;66:1527–42.

31. Zheng C, Acheampong AK, Shi Z et al. Abscisic acid catabolism enhances dormancy release of grapevine buds. Plant Cell Environ 2018;41:2490–503.

32. Rubio S, Pérez FJ. ABA and its signaling pathway are involved in the cold acclimation and deacclimation of grapevine buds. Sci Hortic 2019;256:108565.

33. Rubio S, Noriega X, Pérez FJ. ABA promotes starch synthesis and storage metabolism in dormant grapevine buds. J Plant Physiol 2019;234–235:1–8.

34. Pérez FJ, Noriega X, Rubio S. Hydrogen Peroxide Increases during Endodormancy and Decreases during Budbreak in Grapevine (Vitis vinifera L.) Buds. Antioxidants 2021;10:873.

35. Zhang Y, Dami I. Improving freezing tolerance of ‘Chambourcin’grapevines with exogenous abscisic acid. HortScience 2012;47:1750–7.

36. Bowen P, Shellie KC, Mills L et al. Abscisic acid form, concentration, and application timing influence phenology and bud cold hardiness in Merlot grapevines. Charles MT (ed.). Can J Plant Sci 2016;96:347–59.

37. Wang H, Blakeslee JJ, Jones ML et al. Exogenous abscisic acid enhances physiological, metabolic, and transcriptional cold acclimation responses in greenhouse-grown grapevines. Plant Sci 2020;293:110437.

38. Diddi N, Lai L, Nguyen CH et al. An efficient and scalable synthesis of a persistent abscisic acid analog (+)-tetralone ABA. Org Biomol Chem 2023;21:3014–9.

39. Willwerth J, Abrams S. Improving Cold Hardiness and Delaying Deacclimation Using Long Lasting Abscisic Acid Analogs. Ontario Grape and Wine Research Incorporated, 2019.

40. Gunn A. Investigating Vitis sp. cold stress tolerance with abscisic acid (ABA) analogs. 2023.

41. Londo JP, Johnson LM. Variation in the chilling requirement and budburst rate of wild Vitis species. Environ Exp Bot 2014;106:138–47.

42. Richardson AD, Duigan SP, Berlyn GP. An evaluation of noninvasive methods to estimate foliar chlorophyll content. New Phytol 2002;153:185–94.

43. Parry C, BLONQUIST Jr. JM, Bugbee B. In situ measurement of leaf chlorophyll concentration: analysis of the optical/absolute relationship. Plant Cell Environ 2014;37:2508–20.

44. Yuan Z, Cao Q, Zhang K et al. Optimal Leaf Positions for SPAD Meter Measurement in Rice. Front Plant Sci 2016;7.

45. Mills LJ, Ferguson JC, Keller M. Cold-Hardiness Evaluation of Grapevine Buds and Cane Tissues. Am J Enol Vitic 2006;57:194–200.

46. Pierquet P, Stushnoff C. Relationship of Low Temperature Exotherms to Cold Injury in Vitis Riparia Michx. Am J Enol Vitic 1980;31:1–6.

47. Richardson EA, Seeley SD, Walker DR. A Model for Estimating the Completion of Rest for ‘Redhaven’ and ‘Elberta’ Peach Trees1. HortScience 1974;9:331–2.

48. North MG, Kovaleski AP. Time to budbreak is not enough: cold hardiness evaluation is necessary in dormancy and spring phenology studies. Ann Bot 2023:mcad182.

49. Kovaleski AP, Reisch BI, Londo JP. Deacclimation kinetics as a quantitative phenotype for delineating the dormancy transition and thermal efficiency for budbreak in Vitis species. AoB PLANTS 2018;10.

50. Eichhorn KW, Lorenz DH. Phenological development stages of the grapevine. Nachrichtenblatt Dtsch Pflanzenschutzdienstes 1977;29:119–20.

51. Kovaleski AP, Londo JP. Tempo of gene regulation in wild and cultivated Vitis species shows coordination between cold deacclimation and budbreak. Plant Sci 2019;287:110178.

52. Luedeling E, Kunz A, Blanke MM. Identification of chilling and heat requirements of cherry trees—a statistical approach. Int J Biometeorol 2013;57:679–89.

53. Santos JA, Costa R, Fraga H. New insights into thermal growing conditions of Portuguese grapevine varieties under changing climates. Theor Appl Climatol 2019;135:1215–26.

54. Wang H, Dami IE, Martens H et al. Transcriptomic analysis of grapevine in response to ABA application reveals its diverse regulations during cold acclimation and deacclimation. Fruit Res 2022;2:1–12.

55. Dobin A, Davis CA, Schlesinger F et al. STAR: ultrafast universal RNA-seq aligner. Bioinformatics 2013;29:15–21.

56. Canaguier A, Grimplet J, Di Gaspero G et al. A new version of the grapevine reference genome assembly (12X.v2) and of its annotation (VCost.v3). Genomics Data 2017;14:56–62.

57. Love MI, Huber W, Anders S. Moderated estimation of fold change and dispersion for RNA-seq data with DESeq2. Genome Biol 2014;15:550.

58. Langfelder P, Horvath S. WGCNA: an R package for weighted correlation network analysis. BMC Bioinformatics 2008;9:559.

59. Grimplet J, Cramer GR, Dickerson JA et al. VitisNet: “Omics” Integration through Grapevine Molecular Networks. PLOS ONE 2009;4:e8365.

60. Subramanian A, Tamayo P, Mootha VK et al. Gene set enrichment analysis: A knowledge-based approach for interpreting genome-wide expression profiles. Proc Natl Acad Sci 2005;102:15545–50.

61. Cutler SR, Rodriguez PL, Finkelstein RR et al. Abscisic Acid: Emergence of a Core Signaling Network. Annu Rev Plant Biol 2010;61:651–79.

62. Nyangulu JM, Nelson KM, Rose PA et al. Synthesis and biological activity of tetralone abscisic acid analogues. Org Biomol Chem 2006;4:1400–12.

63. Fennell AY, Schlauch KA, Gouthu S et al. Short day transcriptomic programming during induction of dormancy in grapevine. Front Plant Sci 2015;6, DOI: 10.3389/fpls.2015.00834.

64. Noriega X, Rubio S, Pérez FJ. Cold Acclimation and Deacclimation Processes in Grapevine Buds: Insights into the Regulation of Cod Hardiness and ICE-CBF-COR Gene Expression. J Plant Growth Regul 2024, DOI: 10.1007/s00344-024-11322-x.

65. Londo JP, Kovaleski AP. Integrating cold hardiness and deacclimation resistance demonstrates a conserved response to chilling accumulation in grapevines. 2024:2024.09.28.615590.

66. Kovaleski AP. Woody species do not differ in dormancy progression: Differences in time to budbreak due to forcing and cold hardiness. Proc Natl Acad Sci 2022;119:e2112250119.

67. Johnson K, Lenhard M. Genetic control of plant organ growth. New Phytol 2011;191:319–33.

68. Nishizawa A, Yabuta Y, Shigeoka S. Galactinol and Raffinose Constitute a Novel Function to Protect Plants from Oxidative Damage. Plant Physiol 2008;147:1251–63.

69. Londo JP, Kovaleski AP, Lillis JA. Divergence in the transcriptional landscape between low temperature and freeze shock in cultivated grapevine (Vitis vinifera). Hortic Res 2018;5:1–14.

70. Majee A, Kumari D, Sane VA et al. Novel roles of HSFs and HSPs, other than relating to heat stress, in temperature-mediated flowering. Ann Bot 2023;132:1103–6.

71. Thompson AJ, Mulholland BJ, Jackson AC et al. Regulation and manipulation of ABA biosynthesis in roots. Plant Cell Environ 2007;30:67–78.

72. Liu Y, Jiang D, Yan J et al. ABA-insensitivity of alfalfa (Medicago sativa L.) during seed germination associated with plant drought tolerance. Environ Exp Bot 2022;203:105069.

73. Hwarari D, Guan Y, Ahmad B et al. ICE-CBF-COR Signaling Cascade and Its Regulation in Plants Responding to Cold Stress. Int J Mol Sci 2022;23:1549.

74. Xiao H, Siddiqua M, Braybrook S et al. Three grape CBF/DREB1 genes respond to low temperature, drought and abscisic acid. Plant Cell Environ 2006;29:1410–21.

75. Tillett RL, Wheatley MD, Tattersall EAR et al. The Vitis vinifera C-repeat binding protein 4 (VvCBF4) transcriptional factor enhances freezing tolerance in wine grape: CBF4 freezing tolerance enhancement in wine grape. Plant Biotechnol J 2012;10:105–24.

76. Dong C, Zhang Z, Ren J et al. Stress-responsive gene ICE1 from *Vitis amurensis* increases cold tolerance in tobacco. Plant Physiol Biochem 2013;71:212–7.

77. Li J, Wang L, Zhu W et al. Characterization of two VvICE1 genes isolated from ‘Muscat Hamburg’ grapevine and their effect on the tolerance to abiotic stresses. Sci Hortic 2014;165:266–73.

78. Karimi M, Ebadi A, Mousavi SA et al. Comparison of CBF1, CBF2, CBF3 and CBF4 expression in some grapevine cultivars and species under cold stress. Sci Hortic 2015;197:521–6.

79. Wang H, Kovaleski AP, Londo JP. Physiological and transcriptomic characterization of cold acclimation in endodormant grapevine under different temperature regimes. Physiol Plant 2024;176:e14607.

80. Chen X-Y, Kim J-Y. Callose synthesis in higher plants. Plant Signal Behav 2009;4:489–92.

81. Singh RK, Maurya JP, Azeez A et al. A genetic network mediating the control of bud break in hybrid aspen. Nat Commun 2018;9:4173.

82. Marowa P, Ding A, Kong Y. Expansins: roles in plant growth and potential applications in crop improvement. Plant Cell Rep 2016;35:949–65.

83. Beauvieux R, Wenden B, Dirlewanger E. Bud Dormancy in Perennial Fruit Tree Species: A Pivotal Role for Oxidative Cues. Front Plant Sci 2018;9.

84. Sudawan B, Chang C-S, Chao H et al. Hydrogen cyanamide breaks grapevine bud dormancy in the summer through transient activation of gene expression and accumulation of reactive oxygen and nitrogen species. BMC Plant Biol 2016;16, DOI: 10.1186/s12870-016-0889-y.

85. Liang D, Huang X, Shen Y et al. Hydrogen cyanamide induces grape bud endodormancy release through carbohydrate metabolism and plant hormone signaling. BMC Genomics 2019;20:1–14.

86. Zhang B, Gao Y, Zhang L et al. The plant cell wall: Biosynthesis, construction, and functions. J Integr Plant Biol 2021;63:251–72.

