## Supporting Material 2 will be used for the link to the file on the preprint site for "Tetralone-ABA enhances winter cold acclimation, reduces deacclimation, and delays budbreak in V. vinifera and V. hybrid grapevines"

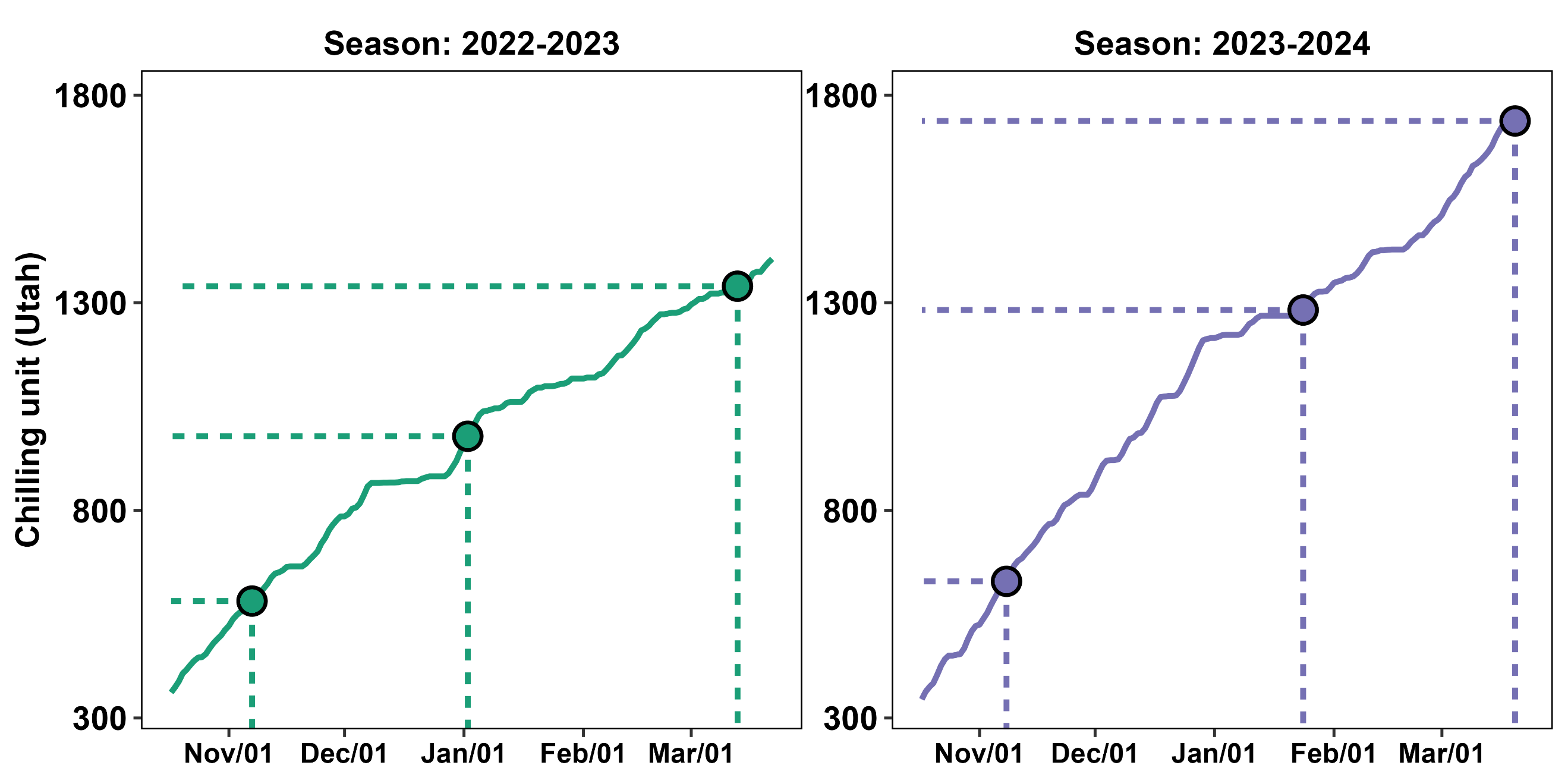

Figure S1. Chilling accumulation (Utah model) in the 2022-2023 and 2023-2024 dormant seasons at the experimental site. Points indicate the timing of field sample collections for the deacclimation assays

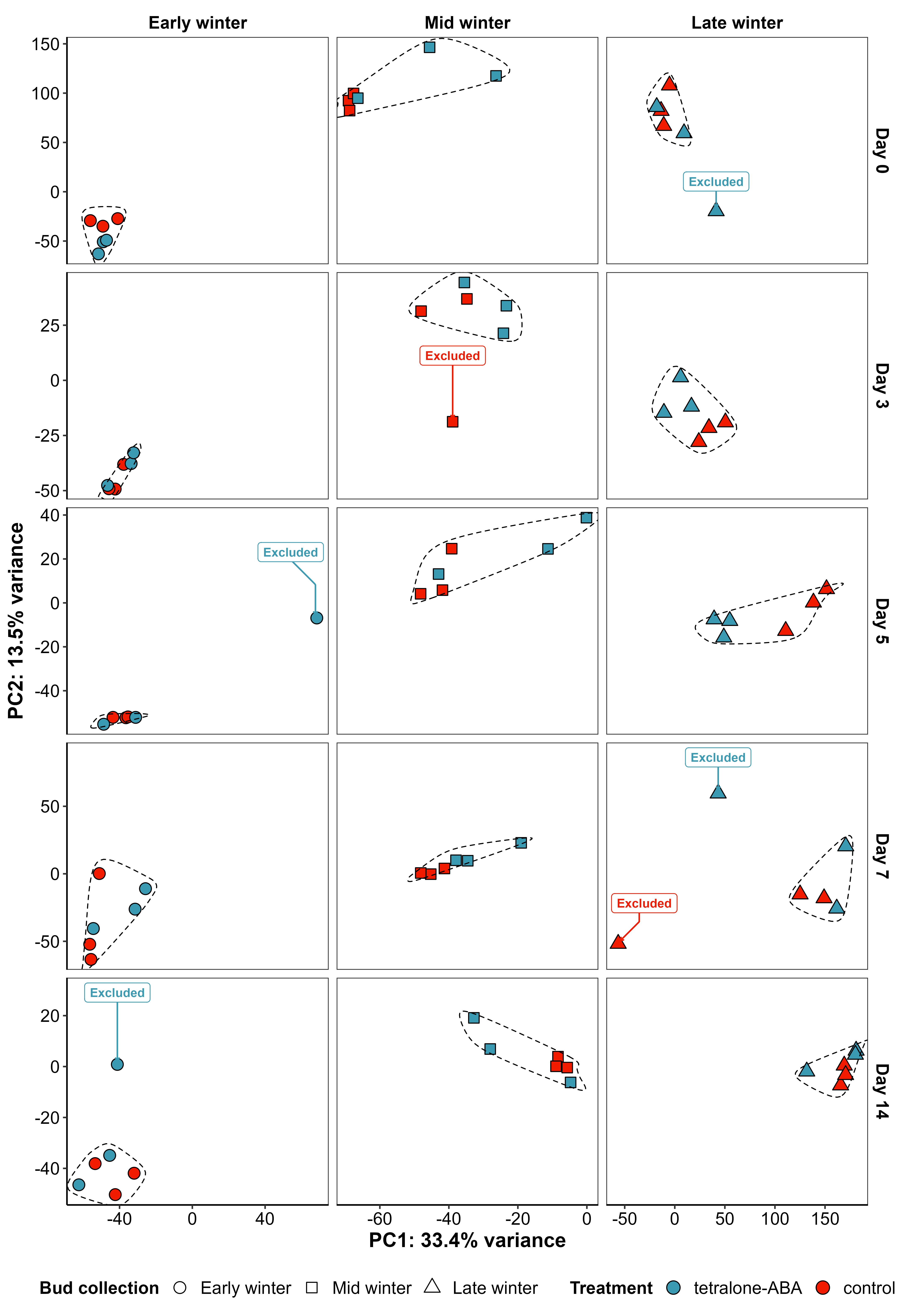

Figure S2. Identification of transcriptome outliers using Principal Component Analysis (PCA)

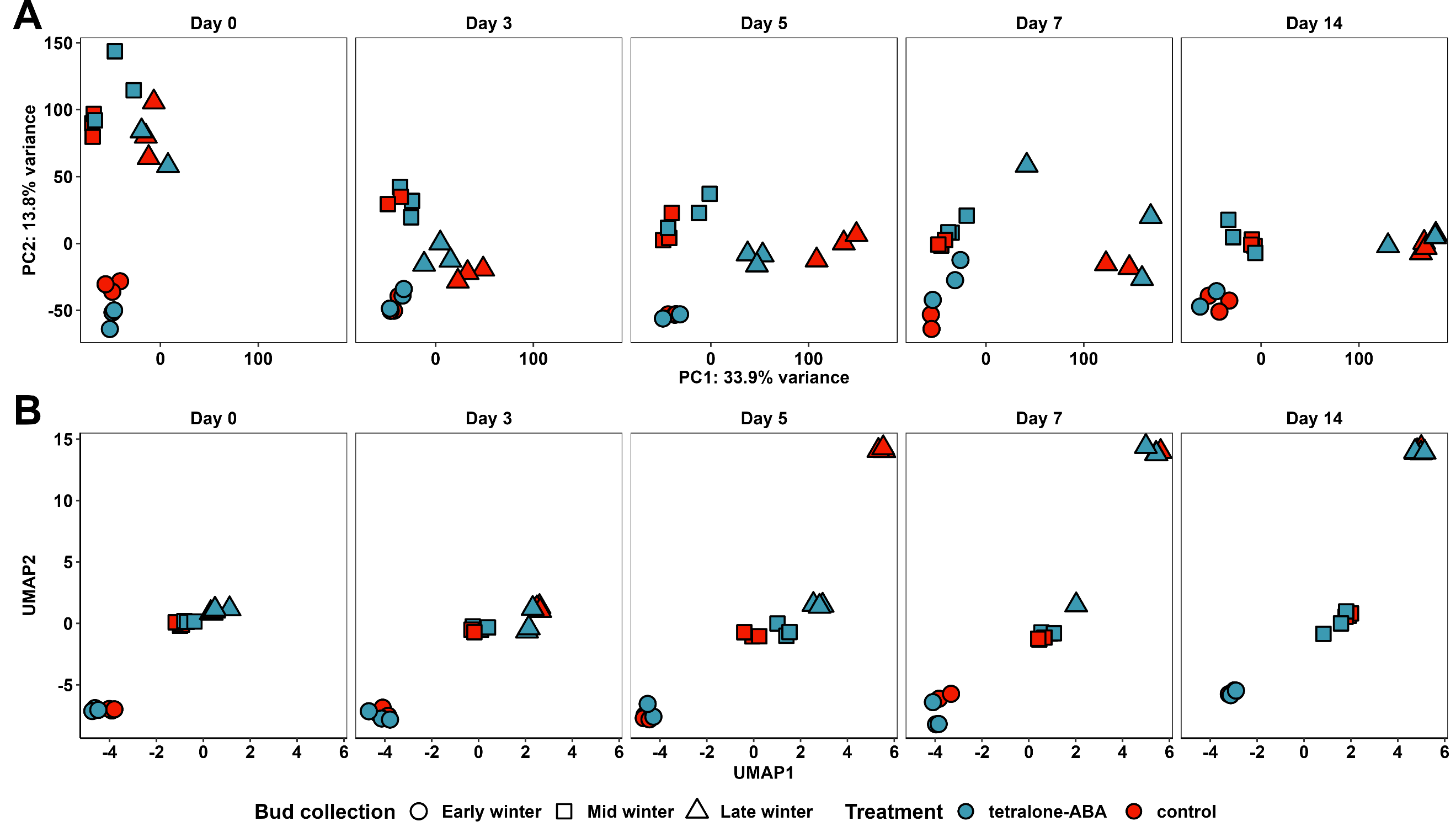

Figure S3. Transcriptome characterization of the RNA-seq libraries. The figure shows the relatedness of all the libraries after outlier filtering and all the genes after low count filtering through A) Principal Component Analysis (PCA) and, B) Uniform Manifold Approximation and Projection (UMAP).

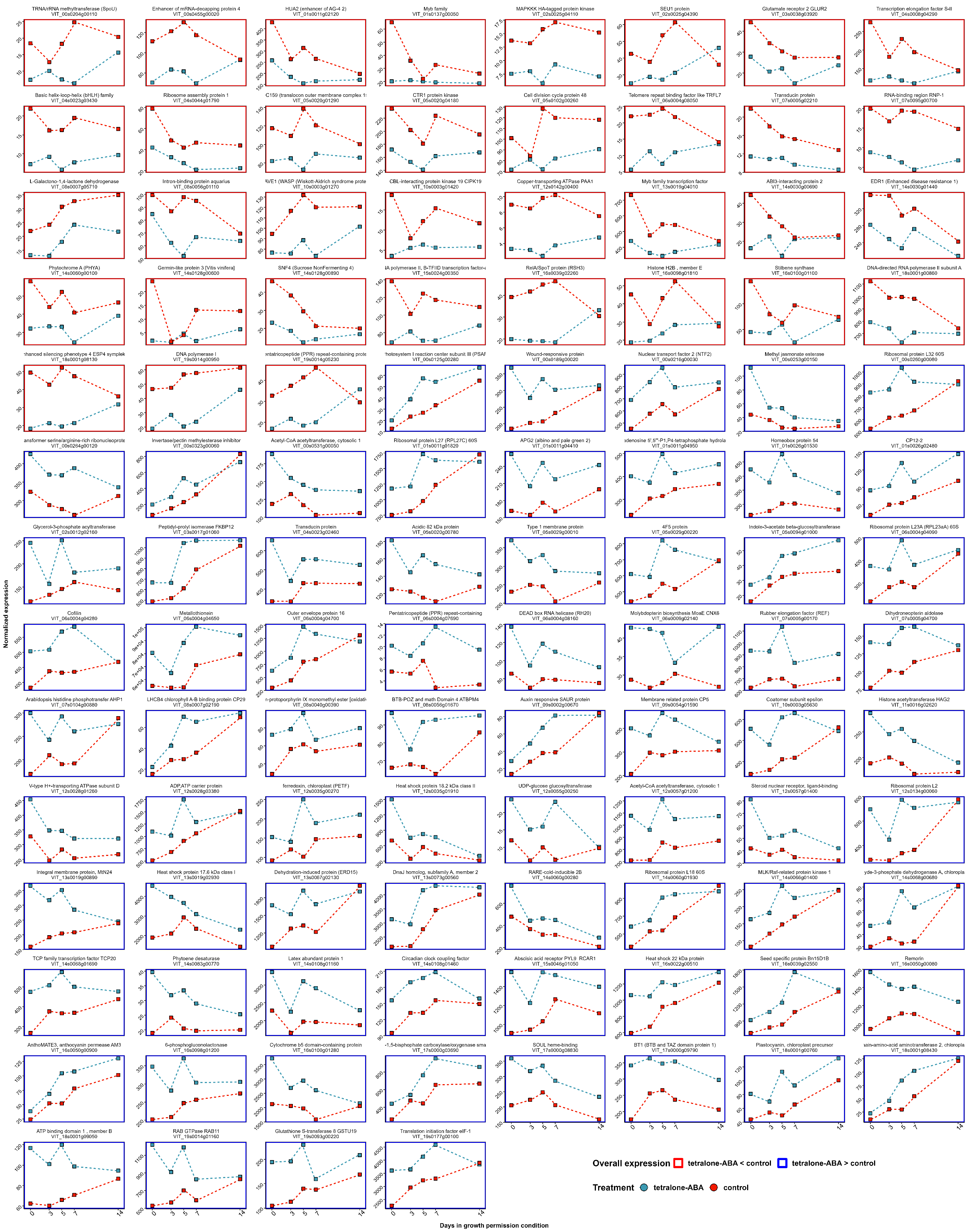

Figure S4. The expression of the differentially expressed genes in the second deacclimation assay. Gene expression shown in the figure represents the mean of DESeq2-normalized expression of all the biological replications.

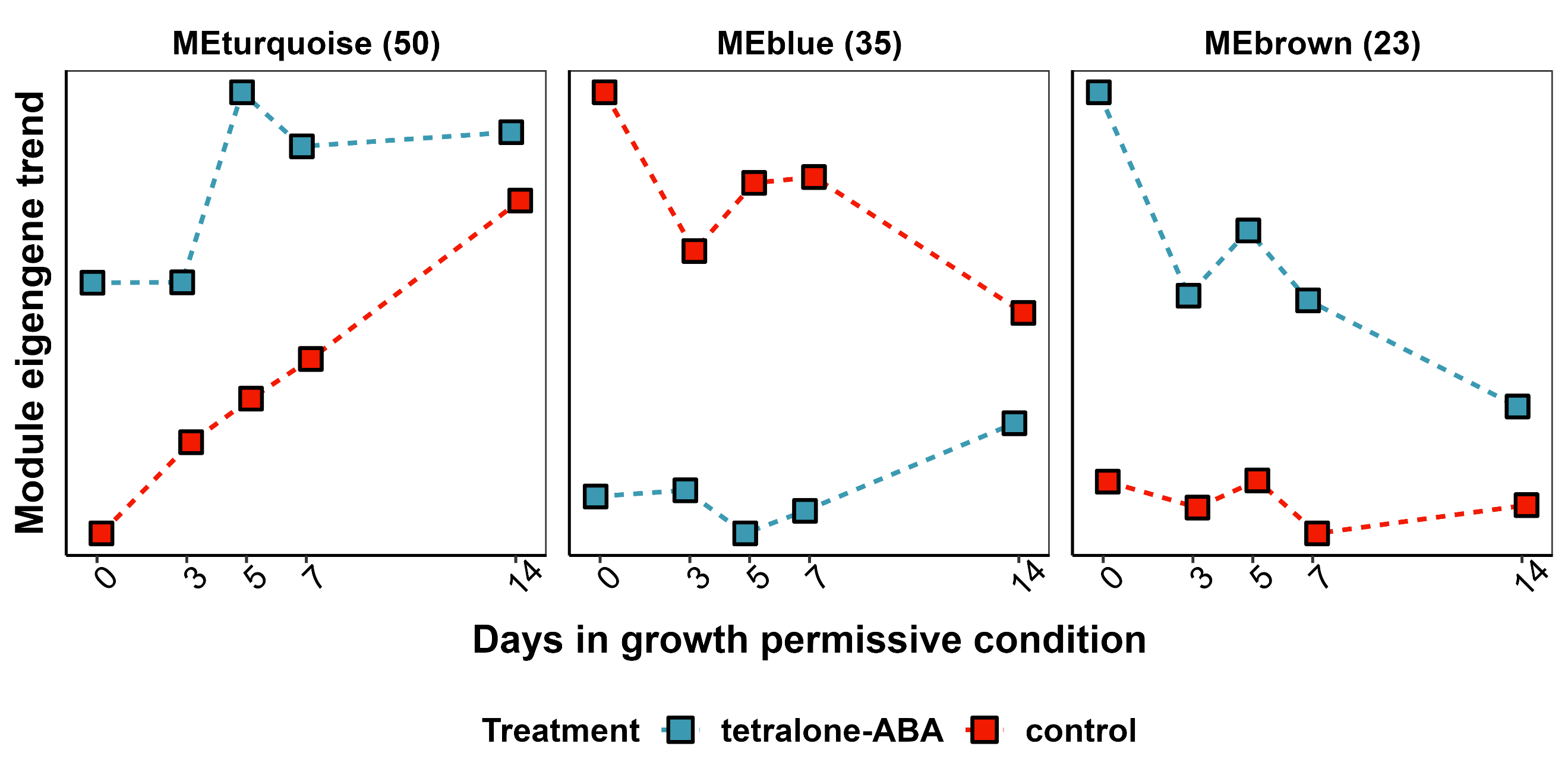

Figure S5. The major expression patterns of the differentially expressed genes in the second deacclimation assay. The expression pattern was identified using weight gene co-expression network analysis (WGCNA).

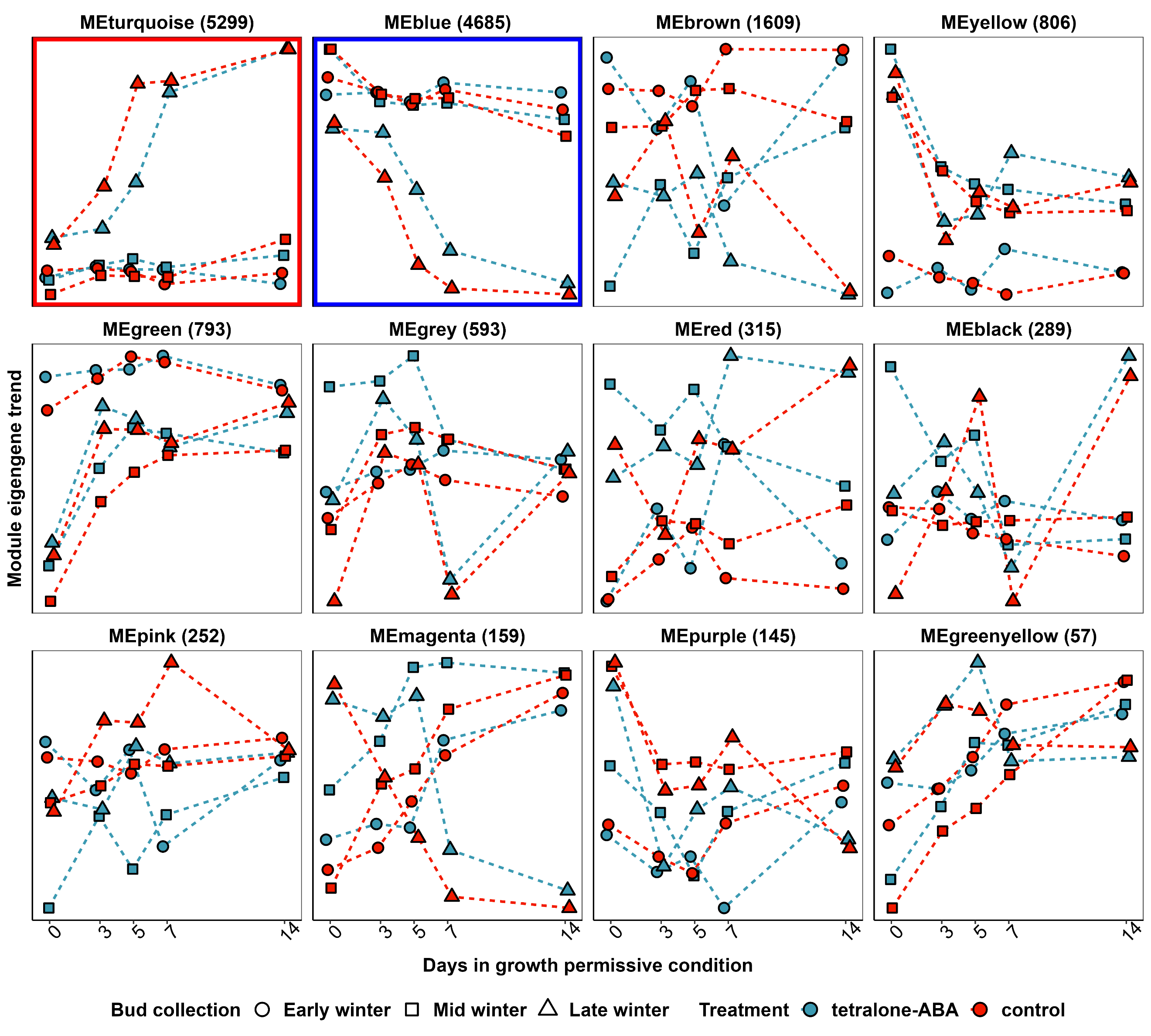

Figure S6. The expression pattern of the module eigengene (ME) of each gene co-expression module identified by weight gene co-expression network analysis (WGCNA) in the deacclimation assays.

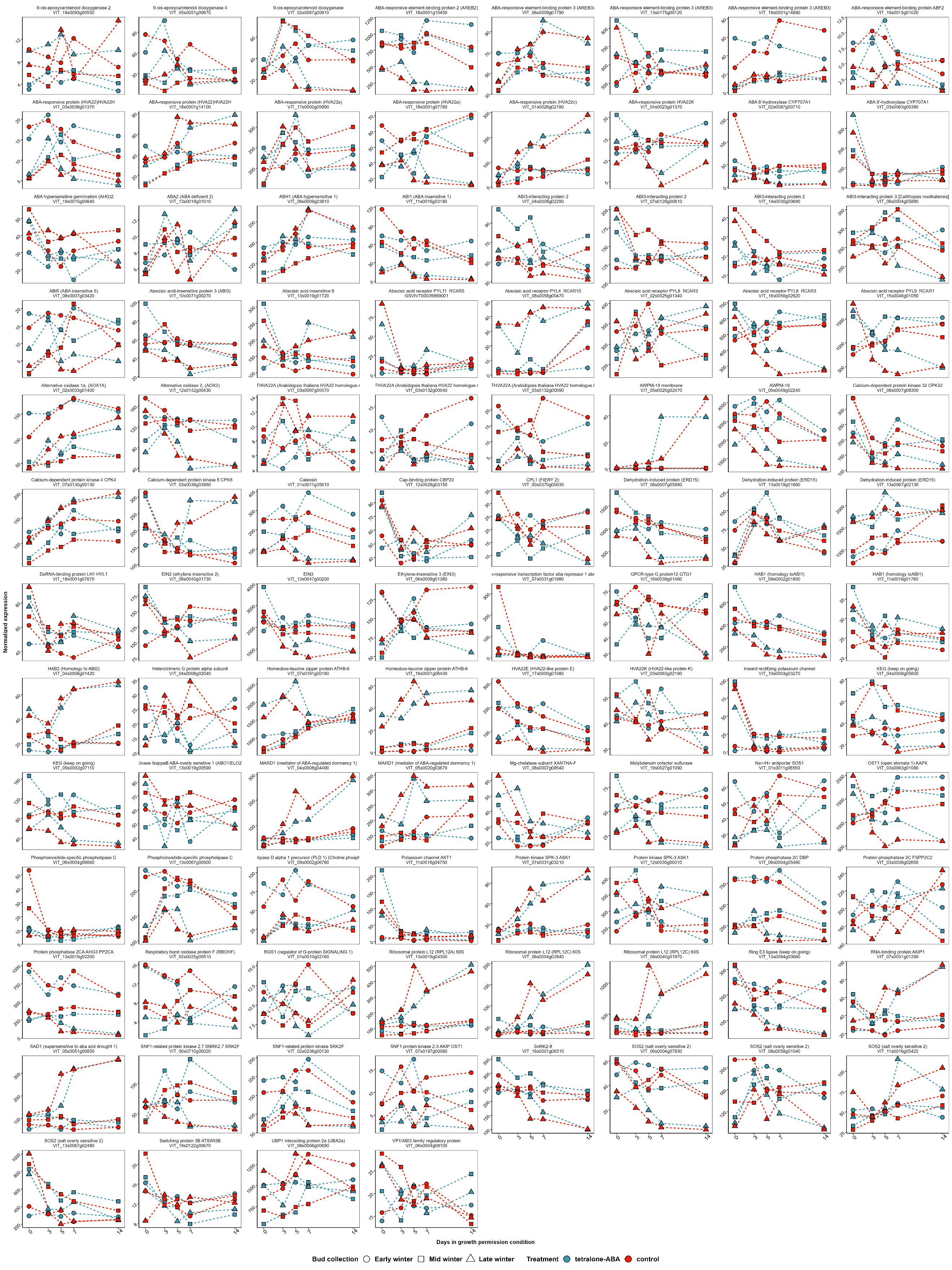

Figure S7. The expression of the genes in ABA biosynthesis and signaling pathways in three deacclimation assays. Gene expression shown in the figure represents the mean of DESeq2-normalized expression of all the biological replications.

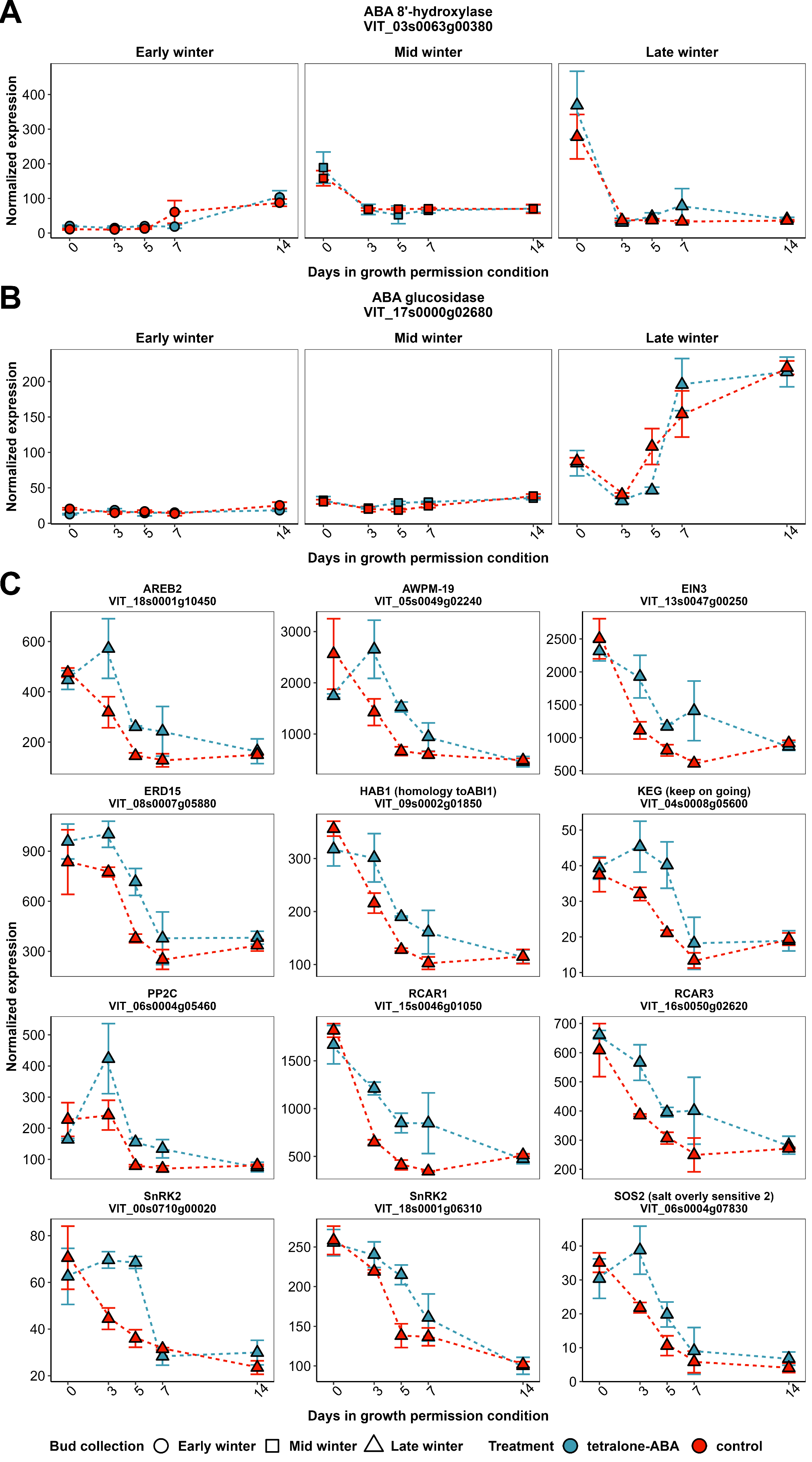

Figure S8. Gene expression pattern of the differentially expressed genes in ABA biosynthesis and signaling pathways. A) The expression of *VIT_03s0063g00380*, that encodes for ABA 8’hydroxylase; B) The expression of *VIT_17s0000g02680*, that encodes for ABA glucosidase; C) The expression of differentially expressed ABA signaling pathway genes and ABA responsive genes in the third deacclimation assays. Gene expression shown in the figure represents the mean of DESeq2-normalized expression of all the biological replications.

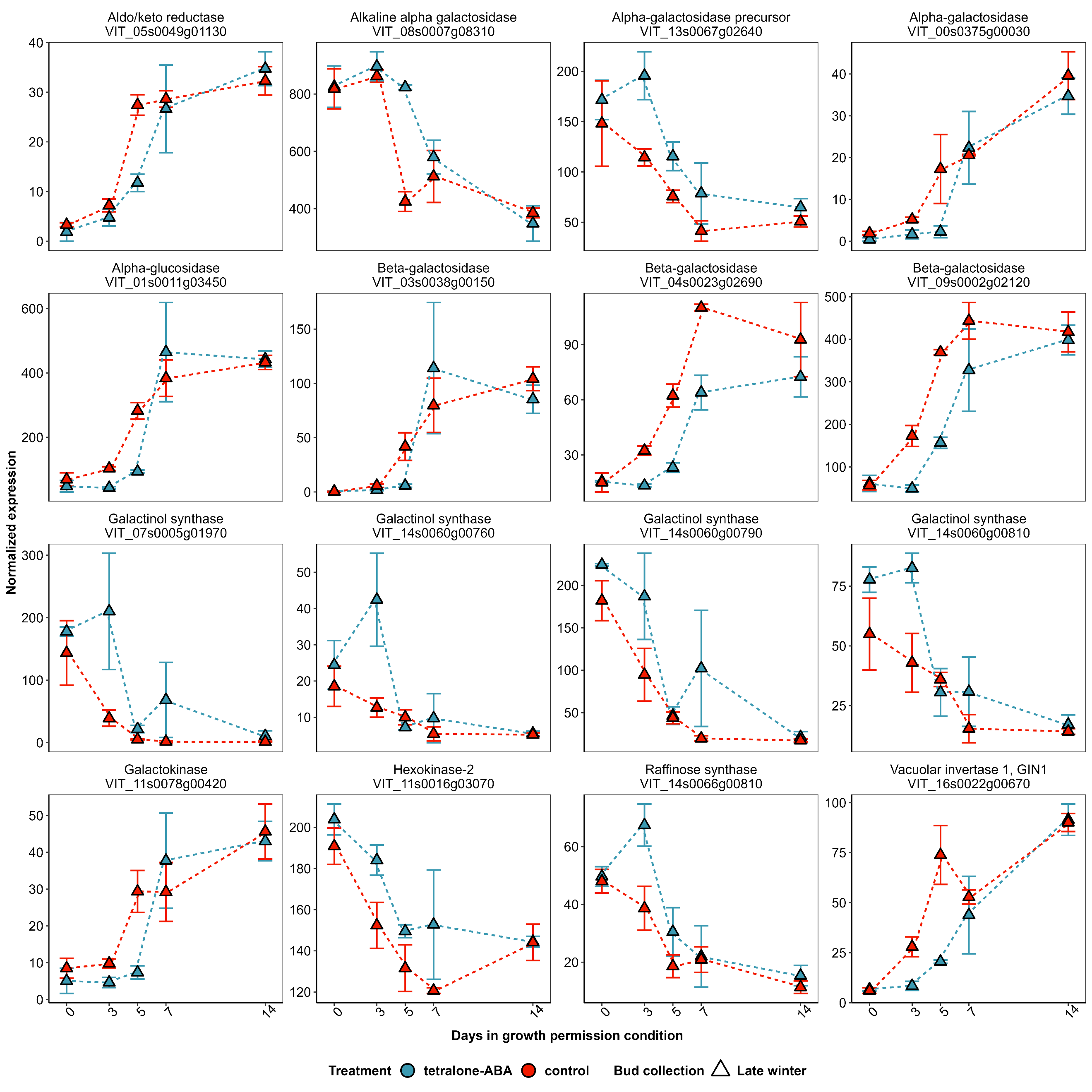

Figure S9. The expression of the genes that showed correlation with deacclimation in galactose metabolism pathway. Gene expression shown in the figure represents the mean of DESeq2-normalized expression of all the biological replications. For clarity, only the libraries collected from the buds in the third deacclimation assay are shown.

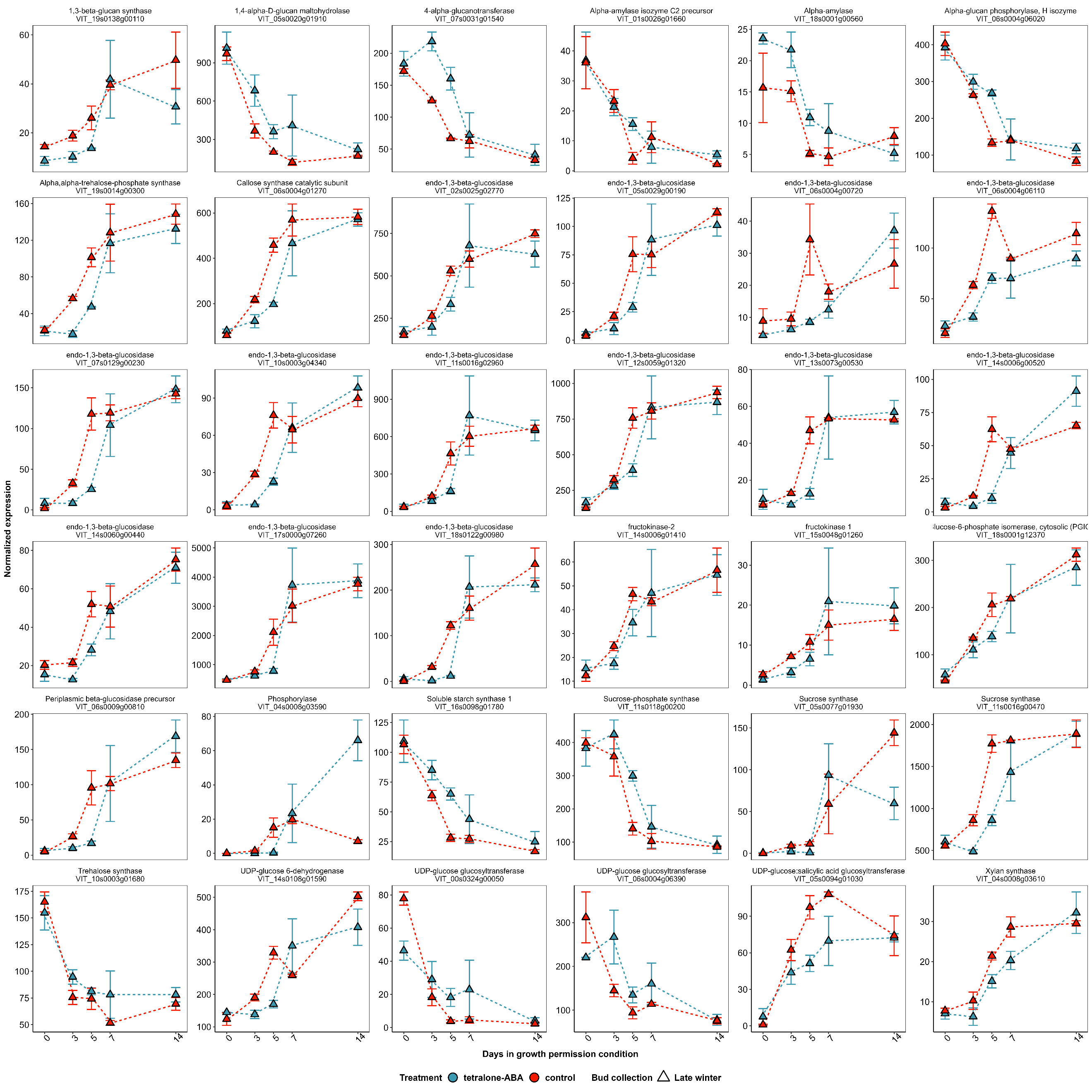

Figure S10. The expression of the genes that showed expression correlation with deacclimation in starch and sucrose metabolism pathways. Gene expression shown in the figure represents the mean of DESeq2-normalized expression of all the biological replications. For clarity, only the libraries collected from the buds in the third deacclimation assay are shown.

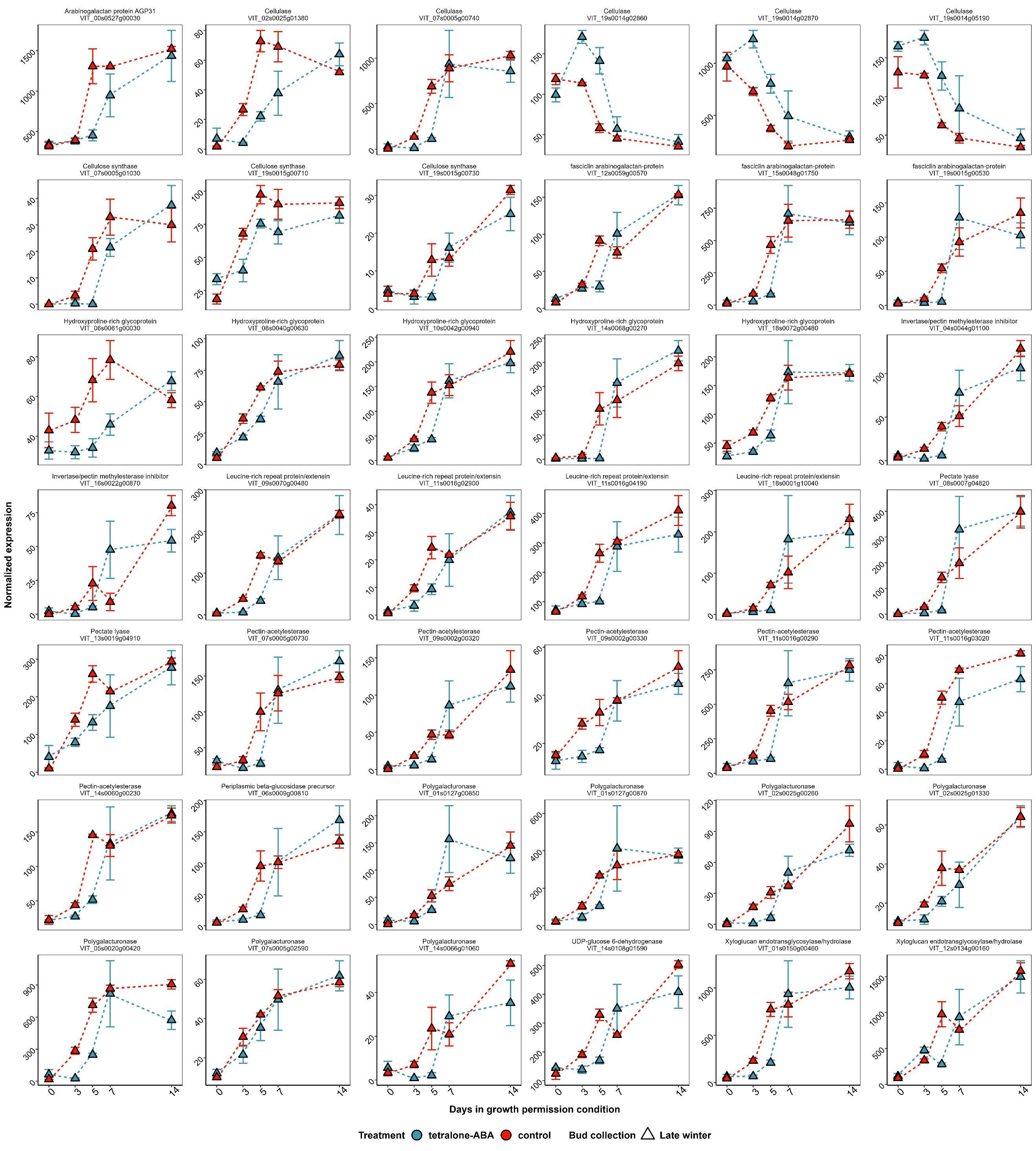

Figure S11. The expression of the genes that showed expression correlation with deacclimation in cell wall pathway. Gene expression shown in the figure represents the mean of DESeq2-normalized expression of all the biological replications. For clarity, only the libraries collected from the buds in the third deacclimation assay are shown.

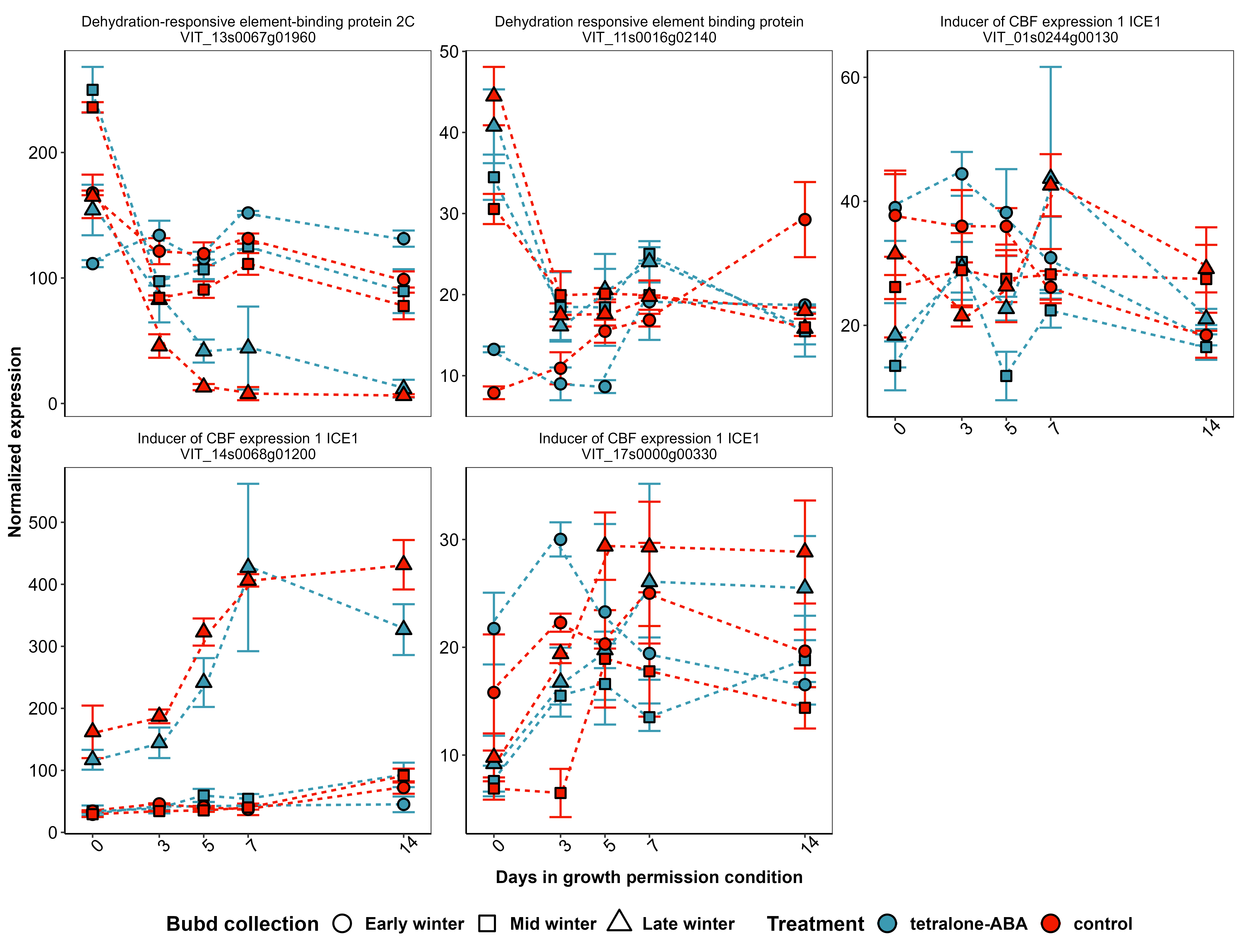

Figure S12. The expression of the ICE, CBF and DREB gene members with detectable expression. Gene expression shown in the figure represents the mean of DESeq2-normalized expression of all the biological replications.

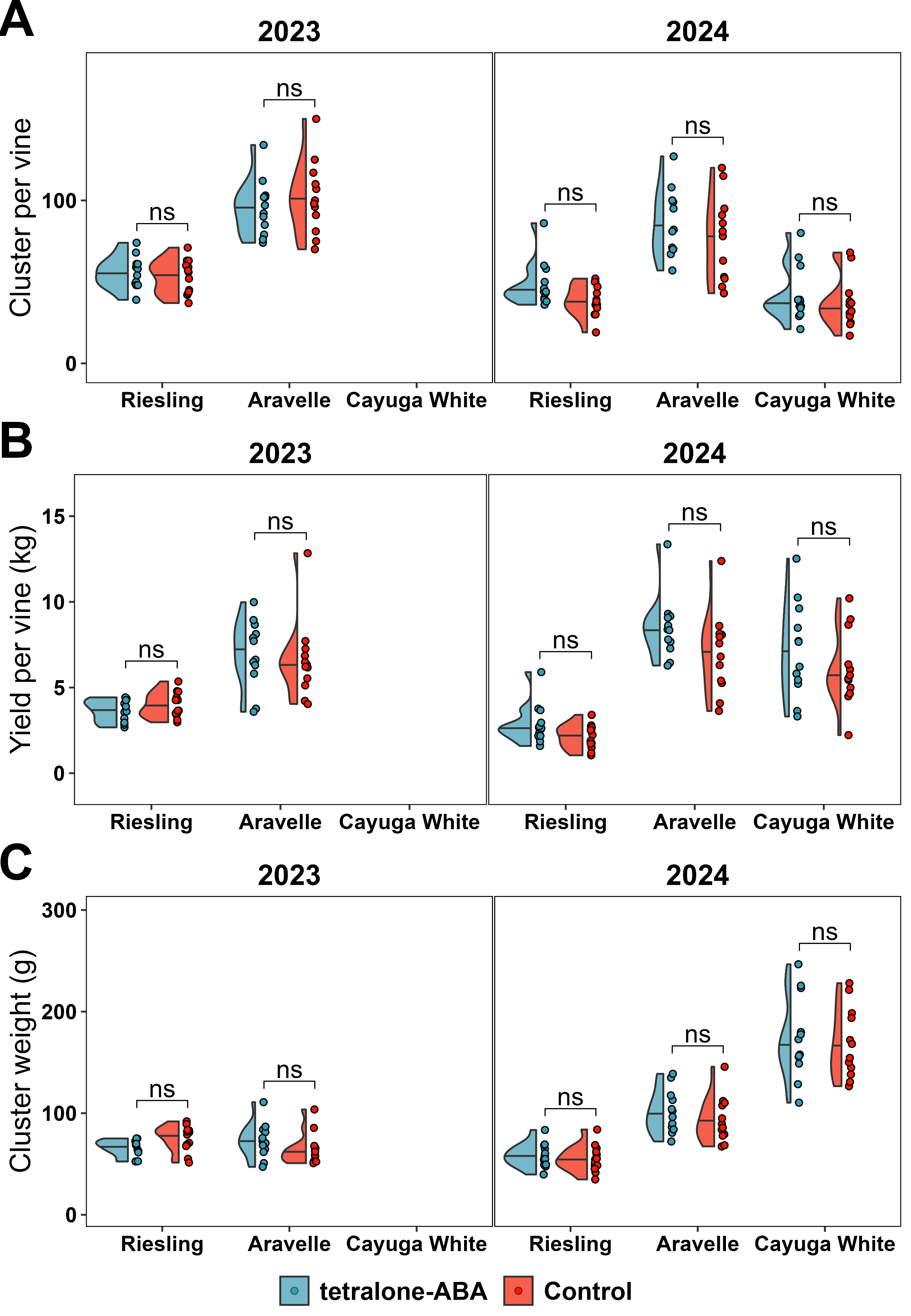

Figure S13. Harvest yield data. A) Cluster per vine; B) Yield per vine; C) Average cluster weight calculated from cluster per vine and yield per vine.

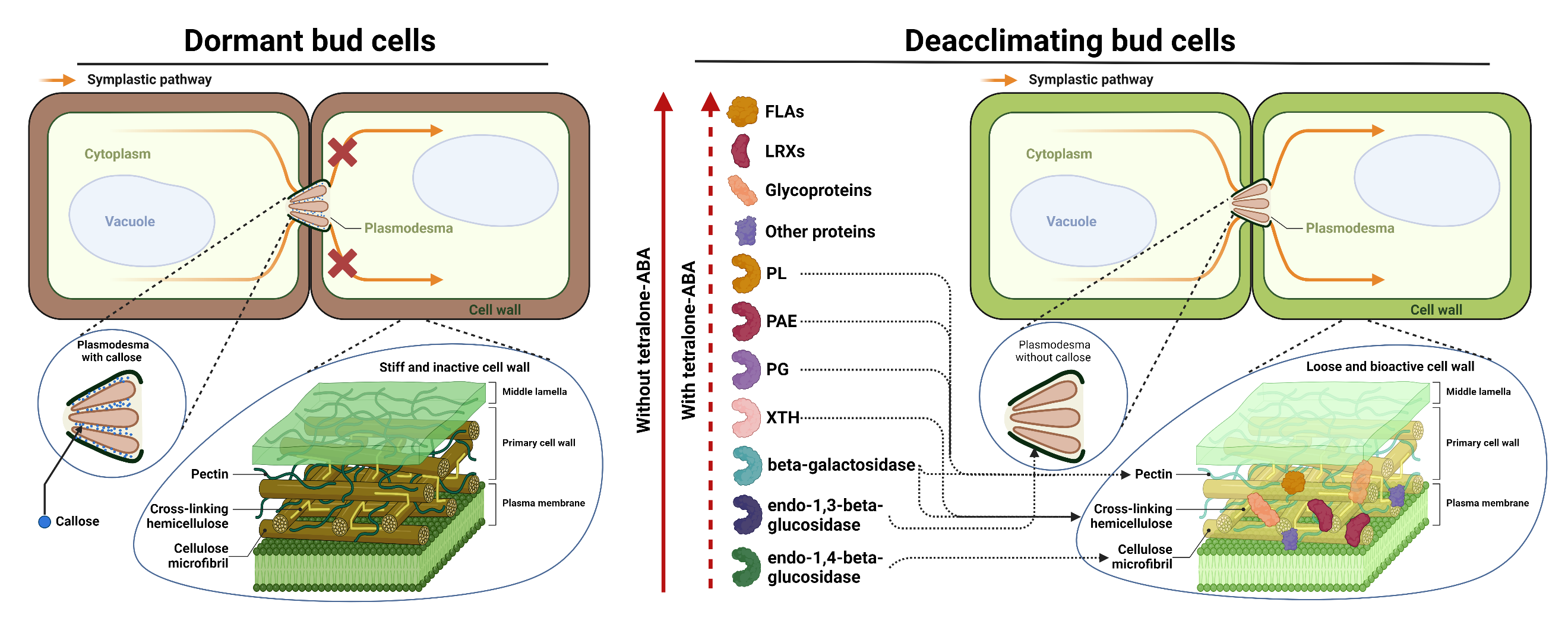

Figure S14. A theoretical cell remodification model during deacclimation and tetralone-ABA’s impact on the model. The line with arrows at the left of the proteins indicates that these proteins were upregulated in control buds during deacclimation. The dashed line indicates the upregulation was delayed in tetralone-ABA treated buds. The lines with arrows linking the proteins and cell wall substrates indicate the proteins’ function in the remodification of the target cell wall substrates.

**Note S1. Functions of proteins encoded by differentially expressed genes in sugar metabolism and cell wall pathways in cell wall modification**

Beta-galactosidases, a diverse family of enzymes in plants, are known for releasing galactose from various substrates like galactolipids, pectin, and xyloglucans, primarily contributing to cell wall hydrolysis and modification and secondary wall deposition [1]. Endo-1,3-beta-glucosidase facilitates the degradation of callose, a type of polysaccharide composed by beta-1,3-glucan with occasional beta-1,6-branches, that is found in the cell walls of higher plant species [2,3]. Some studies revealed that the dynamic build-up of callose is crucial in controlling the bud dormancy process in perennial plants [4,5]. In conditions of short-day photoperiod, ABA in poplar shoot tips leads to an increase in callose production, which results in callose plugs at plasmodesma, leading to the isolation of meristem from symplasmic continuity and promotes dormancy [2,6]. Degradation of callose is proposed to be crucial for the restoration of plant development after dormancy release [7]. FLAs belong to Arabinogalactan-protein family that exhibits in diverse cell wall glycoproteins potentially involved in signaling transduction and plant development such as hormone signaling, cell growth and division, somatic cell embryogenesis, xylem differentiation and response to abiotic stress [8,9]. LRXs are a group of chimeric proteins in the hydroxyproline-rich glycoproteins superfamily that are insoluble in the cell wall and create platforms for protein-protein interactions. They attach to rapid alkalinization factors, influencing cell wall expansion, and engage directly with the FERONIA transmembrane receptor, a key player in regulating cell growth, thus acting as a connection between the cell wall and plasma membrane, detecting signals from the external environment and indirectly transmitting this information to the cytoplasm [10]. PAE, PL, and PG are involved in the modification of pectin, a polysaccharide being the major component of the plant cell wall. PAE does not degrade but modifies the structure of pectin by removing acetyl groups, thus altering its degree of esterification, making it more accessible to degradation by other enzymes. While PL and PG function in the degradation of pectin, PG directly catalyze the hydrolytic cleavage of the alpha-1,4-glycosidic bonds in esterified and de-esterified pectin, but PL uses a beta-elimination mechanism, cleaving the glycosidic bonds in only de-esterified pectin (pectate) [11,12]. XTH is involved in the modification of hemicelluloses, another group of cell wall polysaccharides composed of branched polymers made up of various sugar monomers that contribute to the mechanical strength and flexibility of the cell wall [13]. XTH catalyzes the cleavage and rejoining (transglycosylation) of xyloglucan chains, allowing the restructuring of the xyloglucan network within the cell wall, which enables cell loosening [14,15].

References:

1. Hernández-Nistal J, Martín I, Dopico B *et al.* Coordinated action of β-galactosidases in the cell wall of embryonic axes during chickpea germination and seedling growth. *Plant Biol* 2014;**16**:404–10.

2. Chen X-Y, Kim J-Y. Callose synthesis in higher plants. *Plant Signal Behav* 2009;**4**:489–92.

3. Perrot T, Pauly M, Ramírez V. Emerging Roles of β-Glucanases in Plant Development and Adaptative Responses. *Plants* 2022;**11**:1119.

4. Tylewicz S, Petterle A, Marttila S *et al.* Photoperiodic control of seasonal growth is mediated by ABA acting on cell-cell communication. *Science* 2018;**360**:212–5.

5. Singh RK, Miskolczi P, Maurya JP *et al.* A Tree Ortholog of *SHORT VEGETATIVE PHASE* Floral Repressor Mediates Photoperiodic Control of Bud Dormancy. *Curr Biol* 2019;**29**:128-133.e2.

6. Singh RK, Maurya JP, Azeez A *et al.* A genetic network mediating the control of bud break in hybrid aspen. *Nat Commun* 2018;**9**:4173.

7. Zhao Y, Pan W, Xin Y *et al.* Regulating bulb dormancy release and flowering in lily through chemical modulation of intercellular communication. *Plant Methods* 2023;**19**:136.

8. Ma Y, Yan C, Li H *et al.* Bioinformatics Prediction and Evolution Analysis of Arabinogalactan Proteins in the Plant Kingdom. *Front Plant Sci* 2017;**8**.

9. Liu E, MacMillan CP, Shafee T *et al.* Fasciclin-Like Arabinogalactan-Protein 16 (FLA16) Is Required for Stem Development in Arabidopsis. *Front Plant Sci* 2020;**11**.

10. Herger A, Dünser K, Kleine-Vehn J *et al.* Leucine-Rich Repeat Extensin Proteins and Their Role in Cell Wall Sensing. *Curr Biol* 2019;**29**:R851–8.

11. Shin Y, Chane A, Jung M *et al.* Recent Advances in Understanding the Roles of Pectin as an Active Participant in Plant Signaling Networks. *Plants* 2021;**10**:1712.

12. Jiao X, Li F, Zhao J *et al.* The Preparation and Potential Bioactivities of Modified Pectins: A Review. *Foods* 2023;**12**:1016.

13. Zhang B, Gao Y, Zhang L *et al.* The plant cell wall: Biosynthesis, construction, and functions. *J Integr Plant Biol* 2021;**63**:251–72.

14. Takahashi D, Johnson KL, Hao P *et al.* Cell wall modification by the xyloglucan endotransglucosylase/hydrolase XTH19 influences freezing tolerance after cold and sub-zero acclimation. *Plant Cell Environ* 2021;**44**:915–30.

15. Wu Z, Cui C, Xing A *et al.* Identification and response analysis of xyloglucan endotransglycosylase/hydrolases (XTH) family to fluoride and aluminum treatment in Camellia sinensis. *BMC Genomics* 2021;**22**:761.
